# Single-cell and spatial transcriptomic analyses of gene therapy-associated retinal inflammation in non-human primates

**DOI:** 10.1101/2025.07.31.667874

**Authors:** Célia Sourd, Joel Quinn, Molly C John, Cristina Martinez-Fernandez de la Camara, Lakshanie Wickramasinghe, Moustafa Attar, Hoda Shamsnajafabadi, Ahmed Salman, Sally A Cowley, Calliope Dendrou, Robert E MacLaren, Jasmina Cehajic-Kapetanovic, Kanmin Xue

**Affiliations:** Nuffield Laboratory of Ophthalmology, Nuffield Department of Clinical Neurosciences, University of Oxford, Oxford, UK; Oxford University Hospitals NHS Foundation Trust, Oxford, UK; The Kennedy Institute of Rheumatology, University of Oxford, Oxford, UK; James and Lillian Martin Centre for Stem Cell Research, Sir William Dunn School of Pathology, University of Oxford, Oxford, UK; Great Ormond Street Hospital for Children NHS Foundation Trust, London, UK

**Keywords:** gene therapy, gene therapy-associated uveitis, AAV, single-cell transcriptomic analysis, spatial transcriptomics

## Abstract

Adeno-associated viral (AAV) vectors are rapidly advancing as gene therapies for inherited and common retinal disorders, but gene therapy-associated uveitis (GTAU) limits their broader application.

To investigate the primate ocular immune response, we administered subretinal AAV gene therapy to two non-human primates (NHPs): NHP1 received AAV2-CAG-hRPE65 (voretigene neparvovec) bilaterally at clinical dose; NHP2 received AAV8-GRK1-hRPGRco alongside an analogous mScarlet reporter vector in separate blebs. Longitudinal assessments over three months included multimodal imaging, electroretinography and cytokine profiling, followed by immunohistological, single-cell and spatial transcriptomic analyses of retinal punches.

Both therapies were well-tolerated, with preserved retinal structure and function. Single-cell RNA-sequencing revealed that the AAV8 vector transduced 80% of cones/rods in treated areas, while AAV2 targeted 30% of retinal pigment epithelium (RPE)/rods. Transgene expression did not correlate with apoptotic markers. Persistent immune infiltration (dominated by myeloid and T cells) suggested a type 1 cell-mediated response. Adjunctive intravitreal anti-TNFα (adalimumab) did not appear to mitigate this anti-viral response. Spatial analysis highlighted microglia migration to the subretinal space, consistent with upregulated cytokines (MCP-1/CCL2, IP-10/CXCL10, IL-8/CXCL8, IL-6), which implicate monocytic phagocytes in driving local inflammation.

These findings elucidate the mechanism of GTAU and identify potential therapeutic targets to prevent immune-mediated complications in retinal gene therapy.

## INTRODUCTION

A large number of adeno-associated viral (AAV) vector-mediated retinal gene therapies are in development for the treatment of inherited retinal degenerations (IRDs) and common retinal disorders such as age-related macular degeneration (AMD) ^1^. Voretigene neparvovec (Luxturna, Spark Therapeutics Inc., USA), a subretinally administered AAV serotype 2 vector encoding human *RPE65* under the control of a ubiquitous CAG promoter (AAV2-CAG-*hRPE65*), was the first retinal gene therapy to gain FDA approval in 2017 for the treatment of IRD associated with bi-allelic mutations in *RPE65* ^2^. More recently, clinical trials of another subretinal gene therapy (cotoretigene toliparvovec, Biogen Inc., USA) for X-linked retinitis pigmentosa demonstrated significant improvements in low-luminance visual acuity and retinal sensitivity ^3,4^. This consisted of an AAV8 vector expressing a codon-optimised human retinitis pigmentosa GTPase regulator transgene (*hRPGRco*) under the control of a photoreceptor-specific G protein-coupled receptor kinase 1 (GRK1) promoter. While the pivotal trial for cotoretigene toliparvovec was insufficiently powered to achieve the primary endpoint for clinical approval, a similar vector using an AAV2 capsid variant (AAV2tYF-GRK1-*hRPGRco*, laruparetigene zovaparvovec (laru-zova), Beacon Therapeutics Inc. USA) is currently undergoing randomised, multi-centre phase 3 trial for X-linked retinitis pigmentosa (http://ClinicalTrials.gov ID: NCT04850118) ^5^.

Despite relative immune privilege of the eye associated with the blood-retinal barrier, vector dose-related intraocular inflammation, also termed gene therapy-associated uveitis (GTAU), has been frequently observed in clinical trials and post-approval studies ^3,6–12^. GTAU could present clinically in the form of vitritis, retinitis, cystoid macular oedema or choroiditis, but may also manifest subclinically in the form of subretinal deposits seen on optical coherence tomography (OCT) ^1^. Furthermore, concerns have arisen with the observation of progressive chorioretinal atrophy (CRA) in patients after treatment with voretigene neparvovec, which appears to correlate with AAV dose and may be a result of inflammatory response in the retina ^13,14^. Study in mouse model has indicated the local immunological reaction to subretinal AAV gene therapy to consist of a chronic type 1 cell-mediated response with infiltration of myeloid and T cell in the retina ^15^. However, the immunological responses in rodents may differ significantly from those in humans, thus primate data is vital to provide more direct understanding of the mechanism of GTAU.

In this study, we sought to gain deep insights into the cellular responses to subretinal AAV gene therapy in non-human primates (NHPs) at single-cell resolution using two clinically important AAV vectors: (i) GMP grade AAV2-CAG-*hRPE65* (voretigene neparvovec) and (ii) engineering grade AAV8-GRK1-*hRPGRco*. We also assessed the effects of intravitreal ant-TNFα antibody, adalimumab, as a potential adjunct to prevent intraocular inflammation following gene therapy based on its efficacy in treating non-infectious uveitis ^16–18^. Our results demonstrate that subretinal administration of AAV vectors leads to the activation and migration of microglia to the subretinal space. Monocytic phagocyte derived pro-inflammatory cytokines and chemokines appear to recruit a chronic type 1 cell-mediated anti-viral response characterized by T cells (mainly CD8 effector memory cells) and myeloid cells. While the magnitude of retinal inflammation varied between different vector products and doses, the nature of the immune cell infiltrate remained consistent. Therefore, our data reveal the key immune ligand-receptor signalling pathways which could be targeted for specific immunomodulation to control GTAU.

## MATERIAL AND METHODS

### AAV vectors

The AAV2-CAG-*hRPE65* vector (voretigene neparvovec-rzyl, Luxturna, Spark Therapeutics Inc. USA) was GMP grade vector surplus from approved human retinal gene therapy. The AAV8-GRK1-*hRPGRco* vector was produced at Nationwide Children’s Hospital (Columbus, OH, USA) Vector Core as an ‘engineering grade’ batch for pre-clinical studies ^3^.

The mScarlet-expressing AAV vectors used for NHP subretinal injections (AAV8-GRK1-*mScarlet*) and iPSC-derived microglia experiments (AAV8-CAG-*mScarlet*-WPRE) were produced via transient transfection of adherent HEK293T cells. Confluent HYPERflasks (Corning, Tewksbury, MA, USA) were transfected with the pDG RepCap plasmid (Plasmid Factory, Bielefeld, Germany) of appropriate serotype and the transgene plasmid in TransIT-VirusGEN Transfection Reagent (Mirus bio, Madison, WI, USA). HYPERflasks were harvested 72 hours post transfection and the cell pellet lysed (Lysis buffer 1M Tris, 150mM NaCl, cOmplete Protease Inhibitor (Roche, Basel, Switzerland) in water, pH 8.5). Lysate was subjected to three freeze-thaw cycles and Benzonase (Merck, Darmstadt, Germany) treated. AAV particles were isolated via an ultracentrifugation iodixanol gradient then concentrated and buffer exchanged for phosphate-Buffered Saline (PBS) with an Amicon Ultra 100k filter unit (Millipore, Burlington, MA, USA). Titre was determined by qPCR with primers specific to the transgene (see **Table S1**). Both *mScarlet* vectors preparations were confirmed to be under the limit of detection of endotoxin, and *in vivo* safety was pre-tested by subretinal administration of 1.5×10⁹ vg in mice without observable adverse effects on OCT.

### Animals

Two rhesus macaques (*Macaca Mulatta*) aged 5 years (NHP1) and 6 years (NHP2), obtained from the Medical Research Council (MRC) Centre for Macaques (Harwell Institute, Salisbury, UK), were used in this study. Animals were handled in accordance with The Animals (Scientific Procedures) Act 1986 and UK Home Office regulations. All procedures were performed under general anaesthesia with sevoflurane and Medetomidine, with additional ketamine IV infusion for ERGs. At baseline, 5 mL of blood were collected. cSLO (confocal scanning laser ophthalmoscopy), OCT (optical coherence tomography) and blue-light autofluorescence (AF) imaging were performed with Heidelberg Spectralis HRA (Heidelberg Engineering, Heidelberg, Germany) and analysed with Heidelberg Eye Explorer (HEYEX). Due to the orientation of the camera, all OCT images are inverted such that the inferior retina appear at the top of the images and the superior retina appear at the bottom. Light and dark-adapted baseline ERGs were obtained using a RETeval (LKC Technologies, Gaithersburg, MD, USA).

25-gauge pars plana vitrectomy was performed in all eyes with subretinal injection of AAV vectors using 38G subretinal cannula (MedONE, Sarasota, FL, USA) and foot-pedal-controlled viscous fluid injection (Alcon Constellation, Alcon, Fort Worth, TX, USA), as per human retinal gene therapy surgery ^19^. NHP1 received 1.5x10^11^ vector genomes (vg) of *voretigene neparvovec* via two subretinal injection blebs in both eyes (**Figure 1**). NHP2 received subretinal injections of 1.25x10^11^ vg of AAV8-GRK1-*hRPGRco* and 2.5x10^11^ vg of AAV8-GRK1-*mScarlet* (a ‘shadow’ vector construct expressing a fluorescent reporter) in two separate blebs to each eye. At the end of the procedure, all four eyes of the two NHPs received subtenon injection of 40 mg triamcinolone acetate (Kenalog, Bristol Myers Squibb, Princeton, NJ, USA) and topical dexamethasone/neomycin/polymyxin B (Maxitrol, Southfield, MI, USA) ointment as standard. In addition, both left eyes received intravitreal injection of 1.5 mg of the humanised anti-TNFα antibody, adalimumab (Amgevita, Amgen Limited, Cambridge, UK). No additional drops or systemic immunosuppression was given.

**Figure 1.**
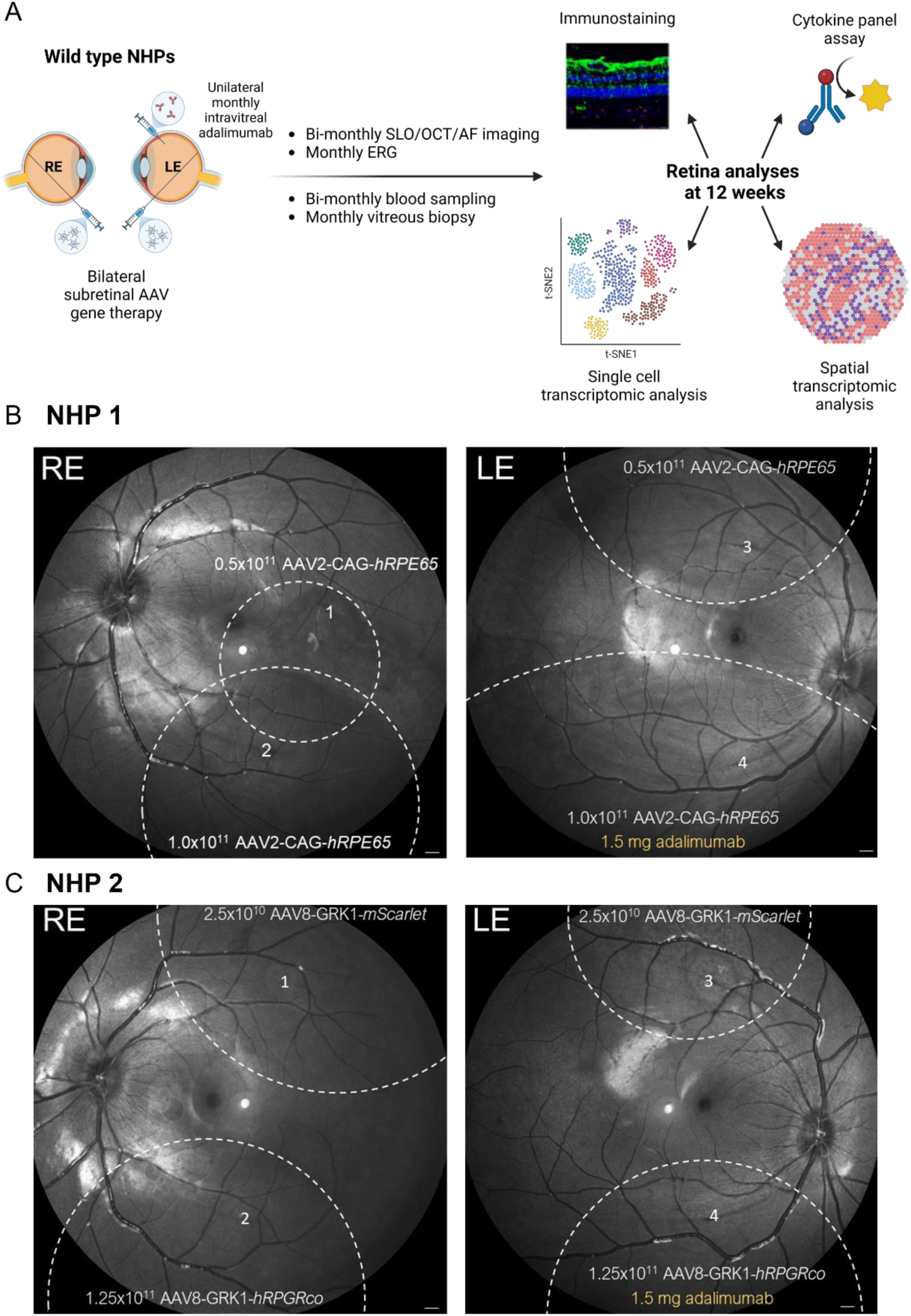
Experimental design. (A) Vitrectomy and subretinal AAV gene therapies were performed to both right (RE) and left (LE) eyes of two wildtype non-human primates (NHPs). The left eyes of both animals also received monthly intravitreal injection of 1.5mg adalimumab. Retinal structure and function were longitudinally monitored by multimodal imaging and ERG. Vitreous and blood samples were collected at key timepoints for cytokine analysis. Retinas were harvested at 12 weeks for immunohistochemistry, single-cell RNA-sequencing, and spatial transcriptomic analysis. (B) NHP1 received a total dose of 1.5x10^11^ vg (vector genome) of Luxturna (AAV2-CAG-*hRPE65*) per eye via two subretinal blebs (denoted by dashed lines). (C) NHP2 received 1.25x10^11^ vg of AAV8-GRK1-*hRPGRco* and 2.5x10^10^ vg of an equivalent reporter vector AAV8-GRK1-*mScarlet* in two separate blebs (dashed lines). Note that all retinal images are vertically inverted (top of the image representing inferior retina) due to the imaging device approaching the supine animal from the top. Scale bars = 500 µm.

Based on the actual vector dose injected, areas of subretinal blebs measured on AF images, and an assumed RPE cell density of 4000 cells/mm^2^ ^20^, we estimated the vector dose per RPE cell for each treated area (**Table 1**). Blood sampling and retinal imaging were performed at baseline (before AAV subretinal injections) and every 2 weeks thereafter. Vitreous biopsy was obtained at baseline then every 4 weeks. Light and dark-adapted ERGs were obtained at baseline, 6 and 10 weeks. Intravitreal injection of 1.5 mg of adalimumab to the left eyes was repeated at 4 weekly intervals. The animals were euthanised, and eyes and spleen were harvested at 12 weeks.

**Table 1.** Summary of subretinal dosing of AAV vectors in two non-human primates (NHPs). Vector dose (vector genome) per RPE cell for each bleb (numbered in accordance with Figure 1B & C) has been estimated based on the actual volume of vector injected, the bleb size, and an estimated RPE cell density of 4000 cells/mm^2^.

| NHP | R/L<br>eye | Subretinal<br>bleb no. | AAV vector | Dose injected<br>(vg) | Bleb area<br>(mm<sup>2</sup>) | Estimated dose per RPE<br>cell |
| --- | --- | --- | --- | --- | --- | --- |
| 1 | RE | 1 | AAV2-CAG-*hRPE65* | 0.5 x 10<sup>11</sup> | 3.4 | 3,676,471 |
|  |  | 2 | AAV2-CAG-*hRPE65* | 1.0 x 10<sup>11</sup> | 28.3 | 883,392 |
|  | LE | 3 | AAV2-CAG-*hRPE65* | 0.5 x 10<sup>11</sup> | 34.9 | 358,166 |
|  |  | 4 | AAV2-CAG-*hRPE65* | 1.0 x 10<sup>11</sup> | 201.1 | 124,316 |
| 2 | RE | 1 | AAV8-GRK1-*mScarlet* | 0.25 x 10<sup>11</sup> | 48.6 | 128,601 |
|  |  | 2 | AAV8-GRK1-*hRPGRco* | 1.25 x 10<sup>11</sup> | 38.5 | 811,688 |
|  | LE | 3 | AAV8-GRK1-*mScarlet* | 0.25 x 10<sup>11</sup> | 22.3 | 280,269 |
|  |  | 4 | AAV8-GRK1-*hRPGRco* | 1.25 x 10<sup>11</sup> | 38.5 | 811,688 |

### Cytokine assays on NHP vitreous and blood samples

NHP serum and vitreous samples were assessed using the LEGENDplex NHP Inflammation Panel with V-bottom Plate (BioLegend, San Diego, CA, USA) according to manufacturer’s instructions. Briefly, serum or vitreous fluid samples were centrifuged at 500xg for 5 min to clear then snap frozen on dry ice for storage at -80°c to be analysed with samples from all timepoints together later. Vitreous samples were all run undiluted, whereas serum samples were diluted 1:4 in assay buffer as per manufacturers recommendations. 25 µL of sample was incubated with pre-defined mixed beads in assay buffer for 2 hours at room temperature with 800 rpm shaking. Samples were spun down at 250xg for 5 min, supernatant discarded, and washed in Wash Buffer twice. Detection Antibodies were added and incubated for 1 hour at room temperature with 800 rpm shaking. SA-PE was added directly to sample wells and incubated for 30 min at room temperature with 800 rpm shaking. Samples were spun down and washed as above, followed by final resuspension in 150 µL Wash Buffer. Samples were ran on the Cytek Aurora’s 5-Laser Full Spectrum Cytometer 16UV-16V-14B-10YG-8R (Fremont, CA, USA), set up as per manufacturer’s instructions to gate for the specified bead populations (https://www.biolegend.com/legendplex). Data was analysed in the LEGENDplex Data Analysis Software (Qognit, BioLegend).

### Dissection and tissue preparation

After rapid harvesting of the globes, the cornea and the lens were first carefully removed. The posterior eyecup was flattened with four diagonal radial cuts and the remaining vitreous carefully removed. The flattened eyecup was transferred to a glass slide. Guided by vascular landmarks on retinal images, 4 mm full-thickness retina/choroid/sclera punches were obtained using skin biopsy punches corresponding to treated and untreated areas and numerically assigned for downstream processing. For single-cell transcriptomic analysis, the retina was lifted away from the underlying RPE/Bruch’s membrane of the biopsy punch for dissociation and library preparation. For spatial transcriptomic analysis, full-thickness punches were embedded in optimal cutting temperature compound (VWR, Radnor, PA, USA) and snap frozen in isopentane for storage (at -80 ͦC) or sectioning as per 10X Genomics protocol. For immunohistochemistry, additional full-thickness punches were incubated in pre-chilled 4% PFA for 30 min at room temperature, then transferred to 1% PFA and stored at 4 ͦC until processing.

### Immunohistochemistry and confocal microscopy

Retinal biopsy punches fixed in 1% PFA were cryosectioned to 10 µm slices at -20 ͦC onto Superfrost plus slides (VWR). Slides were washed with PBS. For sections stained with anti-IBA1 (Fujifilm Wako, Osaka, Japan), an additional antigen retrieval step was performed. Briefly, slides were immersed with a pre-heated solution of 10 mM citrate buffer (Thermo Fisher Scientific, Waltham, MA, USA) and 0.05% Tween 20, then heated several times at high temperature. After cooling the slides at room temperature for 20 min, samples were washed with 0.05% Tween-20 in PBS. All conditions were blocked with 10% Normal Donkey Serum (NDS), 0.05% TritonX100 in PBS. Slides were then incubated overnight at 4 ͦC with primary antibodies (full list in **Table S2**) in a solution composed of 1% NDS, 0.05% Triton X-100 and PBS. The next day, slides were washed with 0.05% Tween-20 in PBS and incubated with secondary antibodies under dark conditions for 2 hours at room temperature. Slides were then briefly washed with 0.05% Tween-20 in PBS before being counterstaining with Hoechst diluted at 1:1000 for 15 min in the dark. Coverslips were mounted in SlowFade® Diamond Antifade Mountant (Thermo Fisher Scientific) and sealed. Z-stack images were captured on a Zeiss LSM 710 confocal microscope and post-analysis was performed in Fiji ^21^.

### Retinal dissociation, single cell library preparation and sequencing

Retinal biopsies were dissociated using the Worthington Papain Dissociation system (Worthington, Lakewood, NJ, USA). Retinas were placed in a solution of 20 U/mL papain, 0.005% DNase I with 1 mM L-cysteine and 5 mM EDTA in Earle’s Balanced Salt Solution (EBSS) for 10 min at 37°C with frequent, gentle agitation. Samples were then diluted by addition of 500 μL of EBSS to inactivate the papain and centrifuged at 300xg for 5 min at room temperature. Pellets were resuspended in 525 μL of a solution containing 1 mg/mL ovomucoid and BSA and 100 U/mL DNase I in EBSS. The resulting suspension was carefully layered over 500 μL ovomucoid/BSA solution and centrifuged at 70xg for 6 min. Supernatant was discarded and cells were resuspended in PBS containing 0.04% BSA.

Library preparation for single cell RNA-sequencing (scRNAseq) analysis of dissociated retinal cells was performed using the Chromium Next GEM 10X Single Cell 5’ v2 Dual Index Kit (10X Genomics, Pleasanton, CA, USA), which includes post GEM-generation clean-up, cDNA amplification and DNA quantification. Libraries were quality controlled and sequenced by Novogene (Cambridge, UK) using the Illumina NovaSeq X (Illumina, San Diego, CA, USA). Transformed raw sequencing data were provided as FASTQ files. The Cellranger MkRef pipeline from 10X Genomics was used to build a custom reference genome using FASTA and GTF files for the Mmul_10 *Macaca mulatta* reference genome appended with additional AAV transgene sequences. Cellranger count was then used to perform alignment, filtering, barcode counting, and UMI counting from FASTQ files to the custom reference genome. This generated feature-barcode matrices for each sample, which was used for downstream analyses.

### Analysis of NHP single-cell RNA-sequencing data

The SoupX package ^22^ was used to correct raw feature-barcode matrices for ambient RNA contamination, followed by doublet detection with scDblFinder ^23^. Corrected matrices were then used for analysis with Scanpy ^24^. Commonly used QC metrics, such as UMI count, number of features, percentage mitochondrial RNA and percentage ribosomal RNA were used to filter out low quality cells, and doublets called by scDblFinder were removed. Samples were then log normalised and highly variable genes were selected. Harmony integration was then performed to remove batch effects and generate a single feature barcode matrix.

The integrated matrix was passed through standard dimensionality reduction and clustering pipelines in Seurat ^25^. Briefly, Principal Component Analysis (PCA) was used to determine dataset dimensionality, followed by shared nearest-neighbour graph construction and dimensionality reduction with the Uniform Manifold Approximation Projection (UMAP) method. Annotated clusters were individually subclustered and iteratively re-processed to further remove low quality droplets and doublets. Differential expression was performed by pseudobulking each identified cell type by sample with decoupleR ^26^ and analysing with the DESeq2 package ^27^. Ranked gene lists from DESeq2 output were used for Gene Set Enrichment Analysis using the Molecular Signature Database (MSigDB) Gene Ontology Biological Process gene sets with the fgsea package in R ^28^. Trajectory inference was performed on myeloid cells with the scFates package ^29^ in Python following the tree analysis guidelines. Prior to trajectory inference, the force-directed graph of myeloid cells was drawn using the “draw_graph” function in Scanpy with the Fruchterman-Reingold algorithm ^30^. The ProjecTILs package ^31^ was used for projecting the T cell subset onto the annotated ProjecTILs dataset using the standard pipeline.

### Spatial transcriptomic library generation, processing and analysis

Following optimization, retinal punch biopsies from NHP1 and NHP2 previously fixed in PFA were sectioned at 10 μm with a cryostat and positioned on a Visium (10X Genomics) spatial gene expression slide. The Visium slide was fixed in pre-chilled methanol at -20°C for 30 min, and stained with Hematoxylin & Eosin (H&E). In parallel, RNA from ten adjacent tissue sections were extracted using RNAse-Free DNase kit (Qiagen, Hilden, Germany) and RNA quality checked using Agilent Tapestation (Agilent Technologies, Santa Clara, CA, USA). The H&E-stained retina sections were imaged with the Zeiss Axioscan 7 microscope slide scanner (Zeiss, Oberkochen, Germany). Following 10X Genomics protocol, retina sections were incubated for permeabilization for 12 min at 37°C and reverse transcription was performed. Next, second strand cDNA was synthetised and total cDNA was denatured. qPCR was performed using 10X Genomics’ specific cDNA primers for cDNA quantification before cDNA amplification and clean-up. The next day, cDNA integrity was measured. Spatial gene expression library was finally constructed and sequenced with Illumina NovaSeq X Plus by Novogene.

FASTQ files containing sequencing reads, previously generated custom Mmul_10 reference genome with additional AAV transgene sequences and TIF image files of H&E-stained retina sections were used as input to the Spaceranger count function. Combined data was analysed with 10X Genomics’ Loupe browser software. The expression of the different genes of interest and their spatial localisation in retina samples were obtained using a LogNorm scale value.

### Differentiation of human iPSC to microglia

Cells were cultured at 37°C in 5% CO_2_. Healthy human iPSCs (https://ebisc.org/BIONi037-A, BIONi037-A, Bioneer, Hørsholm, Denmark) were differentiated into microglia using previously described protocol ^32^. Briefly, batch-QCed iPSCs were plated in Geltrex-coated well plates in mTeSR1 medium (STEMCELL Technologies, Vancouver, Canada) supplemented with Rho kinase inhibitor upon thaw to maintain viability during single-cell suspension (Y27632 Abcam, Cambridge, UK). The cell medium was changed daily, then cells were centrifuged at 400xG into AggreWell 800 plate (STEMCELL technologies) with embryoid body (EB) medium (mTeSR1 medium supplemented with BMP4 at 50 ng/mL (Invitrogen, Thermo Fisher Scientific); VEGF at 50 ng/mL (Invitrogen); SCF at 20 ng/mL (Miltenyi Biotec, Bergisch Gladbach, Germany); and Penicillin-Streptomycin (P/S) at 100X (Gibco, Thermo Fisher Scientific)). After 4 days with daily feeding, EBs were separated from debris with a 40 µM cell strainer and plated in T175 flasks with myeloid differentiation medium: X-VIVO 15 cell media (Lonza, Basel, Switzerland) supplemented with 1X GlutaMax (Gibco, Thermo Fisher Scientific), 1X 2-Mercapto-ethanol (Gibco), M-CSF at 100ng/mL (PHC9501, Invitrogen) and IL-3 at 25 ng/mL (Invitrogen). Differentiation cultures were fed weekly and emergent primitive macrophage precursors were harvested from the supernatant 8 weeks after for microglia differentiation. Primitive macrophage precursors were separated from EBs with a 40 µM cell strainer, counted and resuspended in microglia medium: 1X Advanced DMEM/F12 (Thermo Fisher Scientific) supplemented with 1X Glutamax, M-CSF at 25 ng/mL, GM-CSF at 10 ng/mL (Invitrogen), TGFB1 at 50 ng/mL (Peprotech, Cranbury, NJ, USA), IL-34 at 100 ng/mL (Peprotech) and P/S at 50 U/mL. Macrophage precursors were then plated in 96-well plates and a 50% media change was performed tri-weekly for two weeks. After macrophage precursor differentiation into microglia, cells were collected for assays.

### Phagocytosis assay and cytokine profiling of iPSC-derived microglia

For phagocytosis assays, all iPSC-derived microglia were treated with pHrodo Green Zymosan Bioparticles (Thermo Fisher Scientific) as phagocytic cargo. Control cells were treated with zymosan only. The ‘AAV8-treated 72h prior’ were transduced with an AAV8-CAG-*mScarlet*-WPRE vector at a Multiplicity of Infection (MOI) of 10000 seven days after microglia differentiation. 72 hours after transduction, the media was changed, and cells were washed three times by gently pipetted fresh media on and swirled the plate to remove any residual AAV vector prior to induction of phagocytosis. The ‘AAV8-treated fresh’ were stimulated with AAV8-CAG-*mScarlet*-WPRE present in cell media during the phagocytosis assay only. The ‘lipopolysaccharide (LPS) treated’ cells were included as a positive control. Four replicate wells were included per condition. The phagosome acidification index was determined by fluorescence microscopy as the total amount of pHrodo Green signal above threshold for each time point over the number of cells, and processed by Fiji.

As performed for the NHP vitreous and blood samples, a similar LEGENDplex Cytokine assay was performed on the human iPSC-derived microglia. Cells were plated in 96-well plates in microglia medium. At baseline, AAV-CAG-*mScarlet*-WPRE was added to the cells at a MOI of 10000. In some wells, adalimumab (Amgevita 20mg, Amgen Limited) was also added to the supernatant at a final concentration of 1 µg/mL. Supernatant was then collected at 1, 4, 8, 24, 72 and 96 hours, assessed using the LEGENDplex Human Inflammation Panel (BioLegend) and analysed as previously described.

## RESULTS

### Longitudinal assessment of retinal structure and function following subretinal gene therapy in non-human primates

In order to simulate the clinical impact of AAV vector-mediated gene therapy and associated immune response, two wildtype NHPs underwent 3-port 23-gauge pars plana vitrectomy and subretinal administration of clinical doses of AAV vectors via 38G subretinal cannula in both eyes, as per current human surgical technique (**Figure 1 A**). The first animal (NHP1) received a total dose of 1.5x10^11^ vector genomes (vg) of clinical grade *voretigene neparvovec* (Luxturna) (AAV2-CAG-*hRPE65*), which expresses human *RPE65* transgene from a ubiquitous CAG promoter, in two separate subretinal blebs in each eye (**Figure 1 B**). The second animal (NHP2) received subretinal administration of 1.25x10^11^ vg of an engineering grade AAV8 vector (AAV8-GRK1-*hRPGRco*) expressing codon-optimised human *RPGR* transgene under the photoreceptor-specific GRK1 promoter in each eye (**Figure 1 C**). In addition, NHP2 also received 2.5x10^10^ vg of a laboratory grade AAV8 vector (AAV8-GRK1-*mScarlet*) in a separate bleb which is analogous to the therapeutic vector but expresses a fluorescent reporter, mScarlet. At the end of surgery, each eye received a subtenon injection of 40 mg of triamcinolone. The left eye of each NHP was also given an intravitreal injection of 1.5 mg adalimumab (an anti-TNFα biologic) which was subsequently repeated every 4 weeks. We had hypothesised that anti-TNFα may reduce gene therapy-associated uveitis (GTAU) based on previous clinical report of its efficacy in non-infectious posterior uveitis ^33^.

Vision recovery following gene therapy was uneventful except for transient photophobia which resolved spontaneously in NHP2. Follow-up examinations under anaesthesia were performed at 2 weekly intervals up to 12 weeks, which included (i) dilated fundal examination with retinal imaging (Heidelberg Spectralis cSLO, OCT and autofluorescence), (ii) blood sampling, (iii) vitreous biopsy (every 4 weeks), and (iv) ERG (every 4 weeks). No clinical signs of media opacity, vitreous or anterior chamber cells were seen during follow-up. Localised hypo-autofluorescence (AF) were seen around the retinotomy sites and perimeter of blebs by 2 weeks, which became more diffuse over time (**Figure 2 A**). In the right eye of NHP1 which was treated with AAV2-CAG-*hRPE65*, marked hypo-autofluorescence and RPE mottling developed in the area of Bleb 1, which correlated with mild disruption of the ellipsoid zone (EZ) on OCT. Notably, this bleb was relatively small in size as it connected with the larger Bleb 2, and consequently received the greatest estimated vector dose per RPE cell (**Table 1**) ^20^. In NHP2, the bleb areas treated with laboratory-grade AAV8-GRK1-*mScarlet* became hyperautofluorescent over the course of 4 weeks, consistent with mScarlet protein expression (**Figure 2 B**). In addition, patchy hypo-autofluorescence developed within the mScarlet vector-treated areas, which in the left eye coalesced over 8 weeks into a wedge-shaped area of RPE atrophy inferior to the retinotomy site. These changes correlated with subretinal infiltrates on OCT, which preceded outer retinal atrophy. In contrast, the areas treated with engineering-grade AAV8-GRK1-*hRPGRco* showed minimal autofluorescence changes except at the retinotomy sites, and no significant changes on OCT. Both the *RPE65* and *RPGR* vectors were well-tolerated, while the *mScarlet* vector was associated with imaging evidence of subretinal inflammation leading to outer retinal atrophy. In the left eye of NHP2, no protective effect from adjunctive intravitreal anti-TNFα treatment was seen against subretinal infiltrates associated with the mScarlet vector treatment. The fovea and central retinal thickness (CRT) of all four eyes remained structurally unchanged (**Figure S1**). At 12 weeks post-gene therapy, all eyes were harvested and processed for immunohistochemistry, spatial transcriptomic and single-cell transcriptomic analyses.

**Figure 2.**
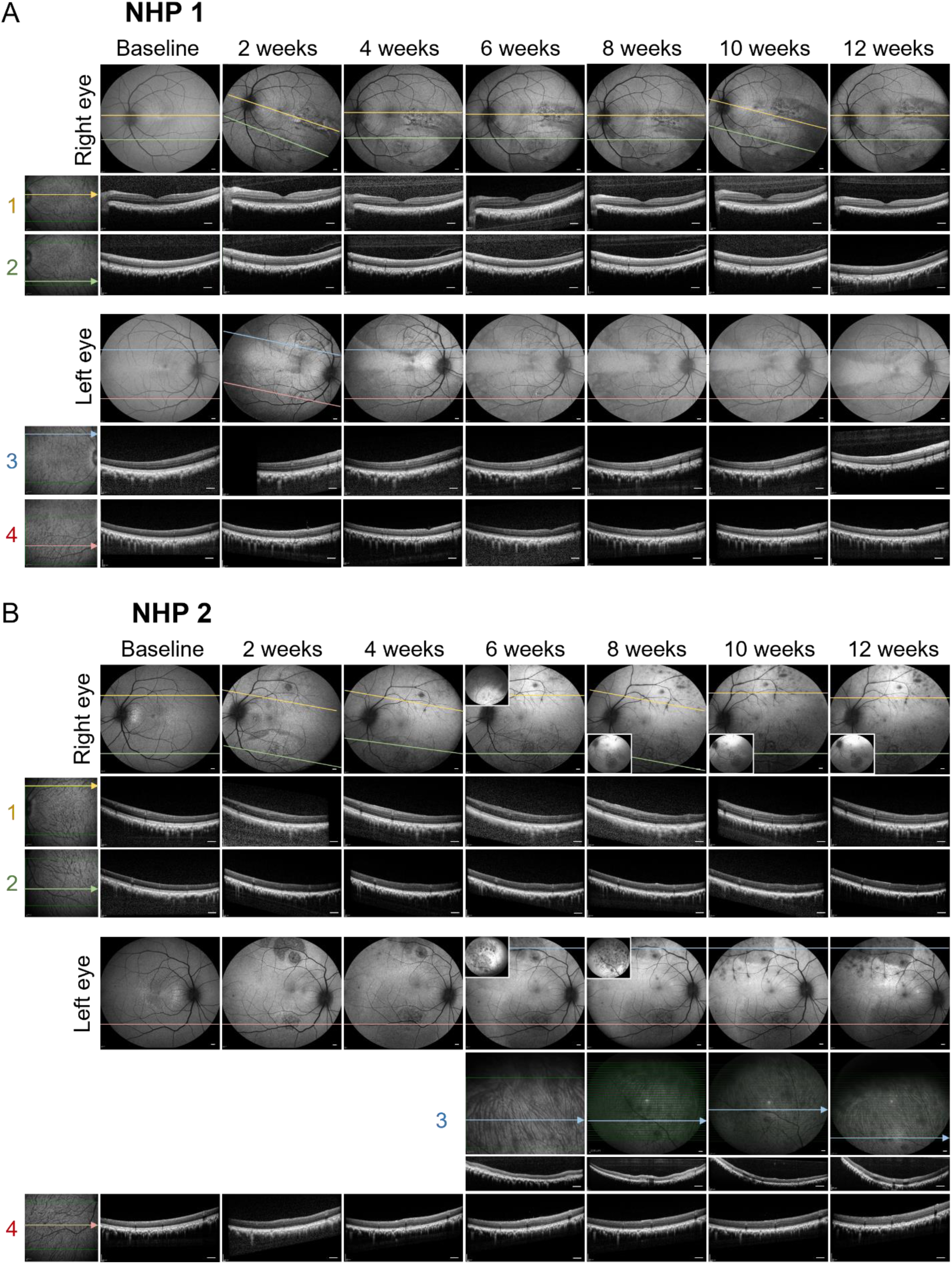
Minimal perturbation of retinal structure and function following subretinal AAV gene therapy in NHPs. Longitudinal multimodal retinal imaging following subretinal injection of AAV vectors over 12 weeks of both eyes NHP1 (A) and NHP2 (B). The colour lines represent the locations of OCT sections from inside the treated subretinal blebs. Note that all retinal images are vertically inverted (top of the image representing inferior retina). Hypo-autofluorescent patches followed by RPE/outer retinal atrophy can be seen along the inferior arcade of NHP2 which was treated with the AAV8.GRK1.*mScarlet* vector. The same area shows increased background hyper-autofluorescence which likely represents mScarlet reporter expression. Scale bars = 500 µm.

Light and dark-adapted ERGs were also obtained at baseline and 10 weeks post-gene therapy. Uveitis has previously been associated with prolonged cone b-wave implicit time (correlated to the peak of the 30Hz flicker response) which could be reversed with treatment ^34,35^. In addition, a change in ERG amplitude of >30% would generally be considered clinically significant ^36^. In NHP1, no significant change in 30Hz flicker implicit time was seen (**Figure S2**). ERG amplitudes were generally unchanged, except for an increase in scotopic b-wave at DA 0.01 cd·s/m² in both eyes (from mean 124 to 208 µV in the right eye and 124 to 178 µV in the left eye). In NHP2, no significant change in 30Hz flicker implicit time was seen (**Figure S3**). However, some reduction in cone response were observed particularly in the right eye: e.g. light-adapted (LA) 30Hz flicker amplitude decreased by 60% (from 113 to 45.3 µV) in the right eye versus 25% (from 120 to 90.3 µV) in left eye; LA b-wave amplitude decreased by 59% (from 118 to 48.7 µV) in the right eye versus 23% (from 121 to 92.6 µV) in the left eye.

### AAV-mediated transgene expression in the retina at single-cell resolution

To evaluate the efficiency of AAV-mediated gene therapy *in vivo*, transgene expression was analysed in both NHP1 and NHP2. Full thickness retinal punches were obtained from areas within and outside the subretinal blebs. Conventional immunohistochemistry was performed on retinal sections (**Figure 3 A and D**). Due to high amino acid sequence homology between macaque and human RPE65 (99%) and RPGR (94%), both the native and transgene-derived proteins were detected with the antibodies. Of note, protein expression appeared higher within the treated blebs than in untreated areas, consistent with vector-mediated transgene expression. Both the native and transgene-derived RPE65 protein appeared to localise to the RPE layer. However, RPGR staining, which is typically localized to the photoreceptor connecting cilia as seen in untreated regions, extended more into the outer nuclear layer (ONL) and outer plexiform layer (OPL) in treated areas (**Figure 3 D**). While such extension of RPGR expression into photoreceptor cell bodies has previously been attributed to fixation artifacts ^37^, it may be a result of AAV-mediated overexpression leading to protein diffusion beyond its normal subcellular localization.

**Figure 3.**
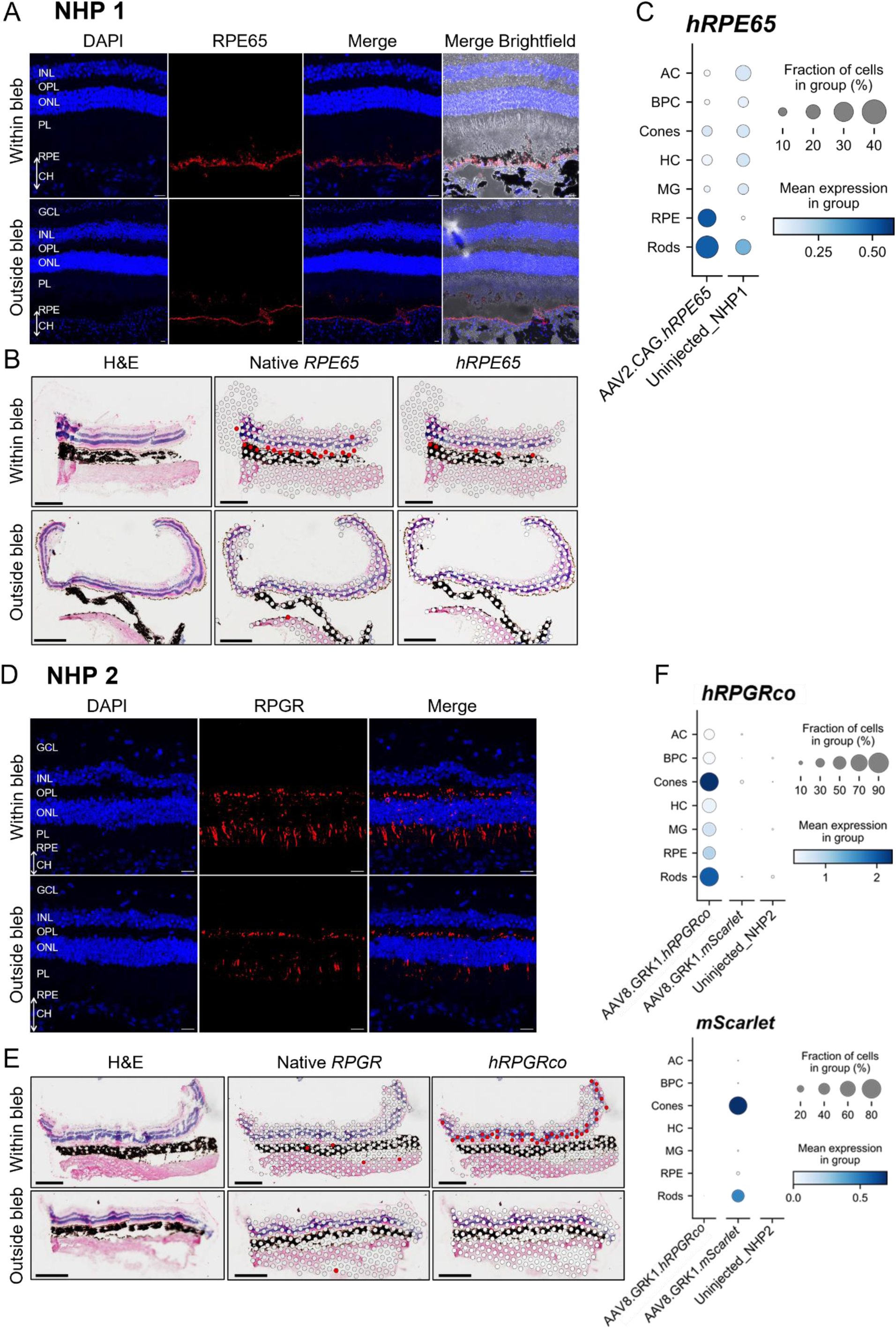
Robust expression of human *RPE65* and *RPGR* transgenes in AAV-treated NHP retinas. (A and D) Immunostaining of retina sections from AAV2-CAG-*hRPE65* (NHP1) and AAV8-GRK1-*hRPGRco* (NHP2) treated areas detect both native NHP and transgene-derived human RPE65 and RPGR proteins, respectively. Control sections were taken from outside the treated blebs. GCL = ganglion cell layer; INL = inner nuclear layer; OPL = outer plexiform layer; ONL = outer nuclear layer; PL = photoreceptor layer; RPE = retinal pigmented epithelium; CH = choroid. Scale bars = 20 µm. (B and E) Spatial transcriptomic maps of native versus human *RPE65* (NHP1) and *RPGR* (NHP2) gene expression within retina sections. Pink spots represent the locations of cell clusters expressing the gene of interest overlayed on the H&E staining image. Scale bar = 0.5 mm. (C and F) Single-cell RNA-seq reveals the levels of transgene expression by each retinal cell type. Circle size represents the proportion of cells that express the transgene within each population, while colour intensity represents the mean expression level per cell. AC = amacrine cells; BPC = bipolar cells; HC = horizontal cells; MG = Müller glia.

Using spatial transcriptomic analysis (Visium, 10X Genomics), we correlated H&E-stained retinal sections with the transcriptome at 55 µm spatial resolution. The technique enabled robust differentiation between native macaque mRNA transcripts and AAV-derived human transgene expression. Human *RPE65* transgene expression can be seen to localise to the RPE layer and expression level appears to be lower than native macaque *RPE65* **(Figure 3 B**). In contrast, human codon-optimised *RPGR* transgene expression was seen at higher level than native *RPGR* in the outer retina (**Figure 3 E**), consistent with the protein overexpression suggested by immunostaining.

Transgene expression was further characterized by single-cell RNA-sequencing (scRNA-seq) of dissociated retina punches taken from within treated bleb areas and untreated control areas of each eye. A total of 89,020 cells were captured following quality control. Clusters corresponding to retinal cell types were identified using marker genes, including amacrine cells (AC; *CALB2, SLC6A9*), bipolar cells (BPC; *TRPM1, GRIK1*), cone photoreceptors (*ARR3, OPN1SW, OPN1LW*), horizontal cells (HC; *ONECUT1*), immune cells (IC; PTPRC), Müller glia (MG; *SLC1A3, GLUL, VIM, CRABP1*), RPE cells (*RDH5, RLBP1, RPE65*), and rod photoreceptors (*NRL, NR2E3, PDE6B, AIF1*) (**Figure S4 A-B**) ^38^. Retinal cell populations were comparable between treated and untreated areas in both NHPs (**Figure S4 C**). In the NHP1 retina treated with AAV2-CAG-*hRPE65*, *hRPE65* transgene expression was detected at high level in approximately 30% of captured RPE cells, at moderate level in 30% of rods, and relatively low level in 15% of cones (**Figure 3 C**). Bipolar cells and Müller glia showed none to minimally detectable level of transgene expression. This distribution appears to reflect the selective tropism of AAV2 for RPE and photoreceptors and efficiency of the ubiquitous CAG promoter among these cell types. In addition, *hRPE65* transgene expression was nearly undetectable outside the subretinal bleb area, indicating minimal lateral spread of the viral vector across the NHP retina. In the NHP2 retina treated with AAV8-GRK1-*hRPGRco*, *hRPGR* transgene expression was detected at high level in 80% of cones, moderate level in 75% of rods, and relatively low level in 40% of RPE cells within the treated areas (**Figure 3 F**). Low level of off-target transgene expression was also detected in approximately 50% of Müller glia, while no significant expression was detected in bipolar cells. Similar distribution and proportion of cellular transduction was mirrored in the areas treated with AAV8-GRK1-*mScarlet* reporter vector, which represents the tropism profile of AAV8. Taken together, these results indicate highly efficient *in vivo* transduction of primate photoreceptors by AAV8 and transgene expression from the photoreceptor-specific GRK1 promoter.

It has been hypothesised that transgene overexpression might cause metabolic stress and cell toxicity. We investigated this by looking for any correlation between the levels of expression of the transgene and apoptotic marker genes at single-cell level using the Apoptosis MSigDB Hallmark gene set among the rod, cone, Müller glia and RPE populations. The expression levels of apoptotic marker genes were found to be low overall and comparable between the treated and untreated areas in both NHPs (**Figure S5**).

### Microglia migration identified by spatial transcriptomic mapping

The retina is considered a relatively immune privileged site due to the presence of the blood-retinal barriers, with microglia being the main resident immune cell population under physiological conditions. Our previous rodent study indicates that subretinal AAV gene therapy induces a delayed cell-mediated immune response in the retina, which is detectable by 2 weeks, peaks around 3-4 weeks and persists significantly beyond ^15^. The timing of this immune response appears to correlate with clinical presentation of GTAU, which is an important limiting factor on treatment safety and efficacy ^11^. The extent of the immune response is known to correlate with vector dose, but may also be affected by the type and preparation of the viral vector ^1,39^.

We analysed retina sections from NHP1 and NHP2 at 3 months post-gene therapy for immune cell populations. In NHP1, immunostaining for the pan-immune cell marker CD45 detected minimal immune cell infiltrate histologically in the *voretigene neparvovec*-treated bleb areas compared with untreated areas (**Figure S6 A**). Staining for GFAP, a marker of gliotic response by Müller cells, did not reveal significant difference between treated and untreated retinas. In contrast, retina sections from NHP2 showed greater CD45 and GFAP staining within vector-treated areas, indicating immune cell infiltration and gliosis (**Figure S6 C**). While immunostaining could detect gross changes in retinal histology, it provides insufficient sensitivity to probe subclinical immune responses. To overcome this limitation, we applied the spatial transcriptomic approach to identify specific immune cell populations through their distinct marker gene expression profiles: e.g. T cells (*CD8A, CD3E, CD4, FOXP3*), B cells (*CD19, CD79A, MS4A1*) and natural killer (NK) cells (*NKG7, NCR1*). No visual differences can be observed when comparing B cell and NK cell population between within and outside the treated bleb from NHP1 (**Figure S6 B**), but NHP2 retina sections displayed a small number of B cell and NK cell clusters in the choroid and the RPE (**Figure S6 D**). In contrast, T cell clusters appeared more abundant inside the AAV-treated blebs of both NHP1 and NHP2 (6-fold change compared to outside the treated bleb).

Furthermore, we looked for evidence of microglia and myeloid cell activity within the retina (**Figure 4**). Visium spatial transcriptomic mapping shows clusters of monocytic phagocytes (expressing marker genes *P2RY12, HEXB and CD68*). The most striking observation was that while the monocytic phagocytes were distributed across different retinal layers in the untreated areas (thus likely to represent resident microglia), they became concentrated in the subretinal space in the treated retinal sections of both NHPs (**Figure 4 A and C**). This is potentially indicative of microglia activation/migration or myeloid cell infiltration to the subretinal space in response to subretinal AAV administration. To corroborate the spatial transcriptomic findings, immunostaining with IBA1 antibody was performed on adjacent retinal sections from the same tissue blocks. While IBA1 staining was very limited in NHP1, IBA1+ cells in NHP2 could be seen to display ramified morphology within the inner and outer plexiform layers in the untreated and treated area (consistent with resting microglia), but more amoeboid-shaped IBA1+ cells in the subretinal space in the treated area (**Figure 4 B and D**).

**Figure 4.**
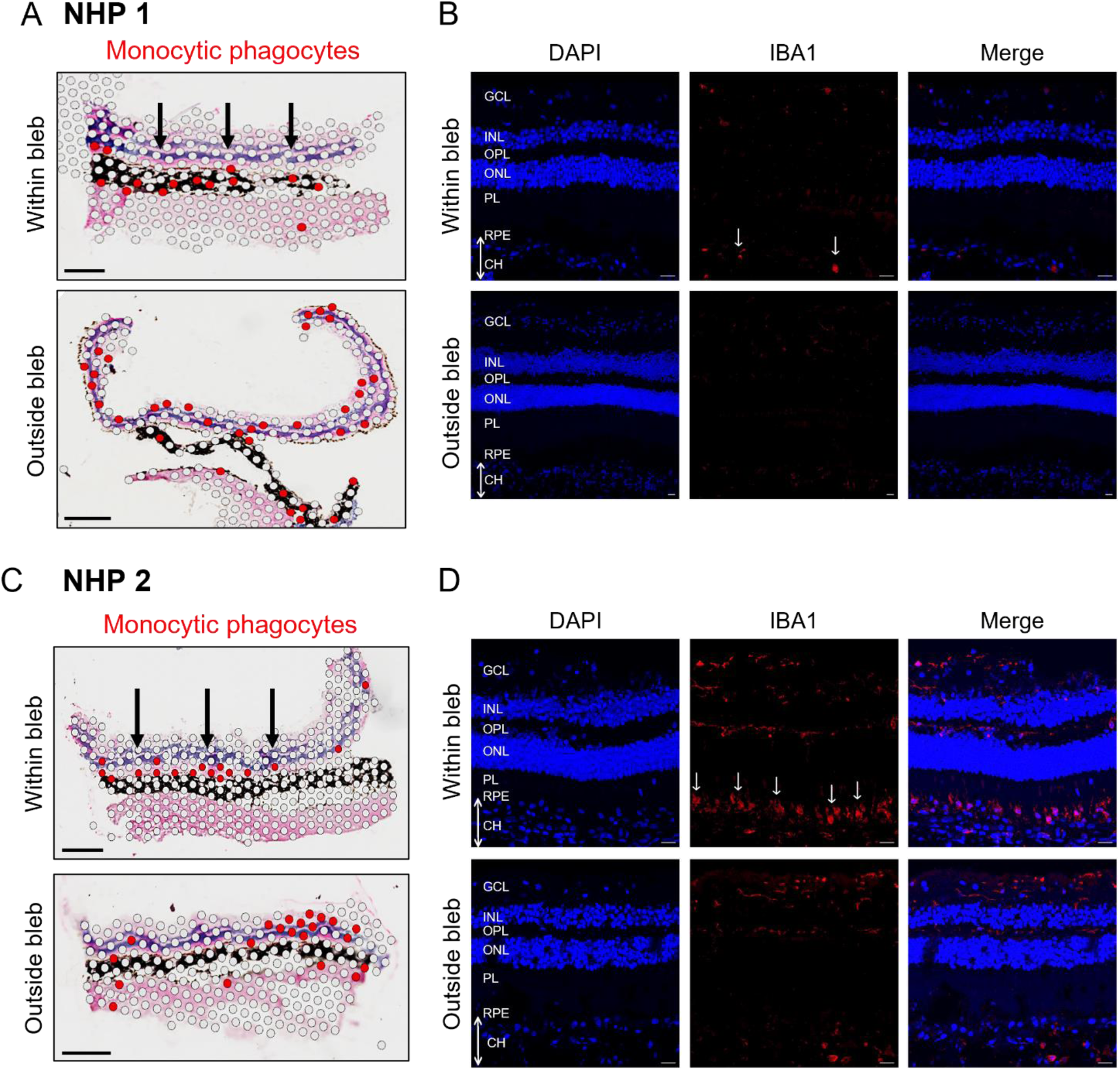
Microglia activation and migration to the subretinal space after AAV gene therapy in NHPs. (A & C) Spatial transcriptomic mapping of monocytic phagocytes (including microglia) over treated versus untreated retinal sections from NHP1 (A) and NHP2 (C). Pink spots represent cells expressing the marker genes P2RY12, HEXB and CD68. Black arrows indicate migration of microglia to the subretinal space in AAV-treated retinas. Scale bar = 0.5 mm. (B & D) Immunostaining for IBA1 demonstrates change in microglia morphology from dendritic to amoeboid, consistent with microglia activation and migration. Scale bar = 20 µm.

### Single-cell characterisation of the nature of gene therapy-associated uveitis (GTAU)

To characterise the nature of retinal inflammation and outer retinal atrophy seen in the AAV8-GRK1-*mScarlet* vector treated region in NHP2, we compared the single-cell transcriptome of retina from this area with an untreated control area from the same eye (**Figure S7**). Differentially upregulated genes were predominantly identified in the rods within the vector-treated area, including *B2M* (MHC I light chain) and *ENSMMUG00000054038* (macaque MHC I antigen), *ENSMMUG00000064120* (ortholog to *CD1D*, MHC-like lipid antigen presenter), ENSMMUG00000050829 (MHC I pathway regulator), *ENSMMUG00000052293* (ortholog to *KIAA1109*, MHC I complex assembly) and ENSMMUG00000058325 (MHC I antigen), which indicate upregulated MHC class I antigen presentation (**Figure S7 A**). A similar set of MHC Class I genes were also upregulated in cones. Moreover, Gene Ontology (GO) enrichment analysis revealed upregulation of gene sets in vector-treated rods that are linked to immune response against viral or cytokine stimuli (**Figure S7 B**). Additionally, of the 58 genes differentially expressed by rods in the AAV8-GRK1-*hRPGRco* bleb, 54 overlapped with the differentially expressed genes in the AAV8-GRK1-*mScarlet* bleb (**Figure S7 C**).

Next, we analysed the immune cell (*PTPRC*^+^) clusters from the whole scRNA-seq dataset, comprising both NHP1 and NHP2. Myeloid cells (*ITGAM*), T/NK cells (*CD3D*) and B cells (*MS4A1*) made up the bulk of the retinal immune cell infiltrate **(Figure 5 A** and **Figure S8**). Interestingly, a cluster of endothelial/smooth muscle cells (*RGS5*) was also present among the annotated immune cell cluster. Myeloid cells were present in significant numbers in all retinal punch areas, including those that were untreated (**Figure S8 B-C**). As expected, the AAV8-GRK1-*mScarlet* and AAV8-GRK1-*hRPGRco* vector-treated areas contributed high proportions of the T cell and B cell clusters, which is indicative of localised adaptive immune response to AAV gene therapy (**Figure S8 C**).

**Figure 5.**
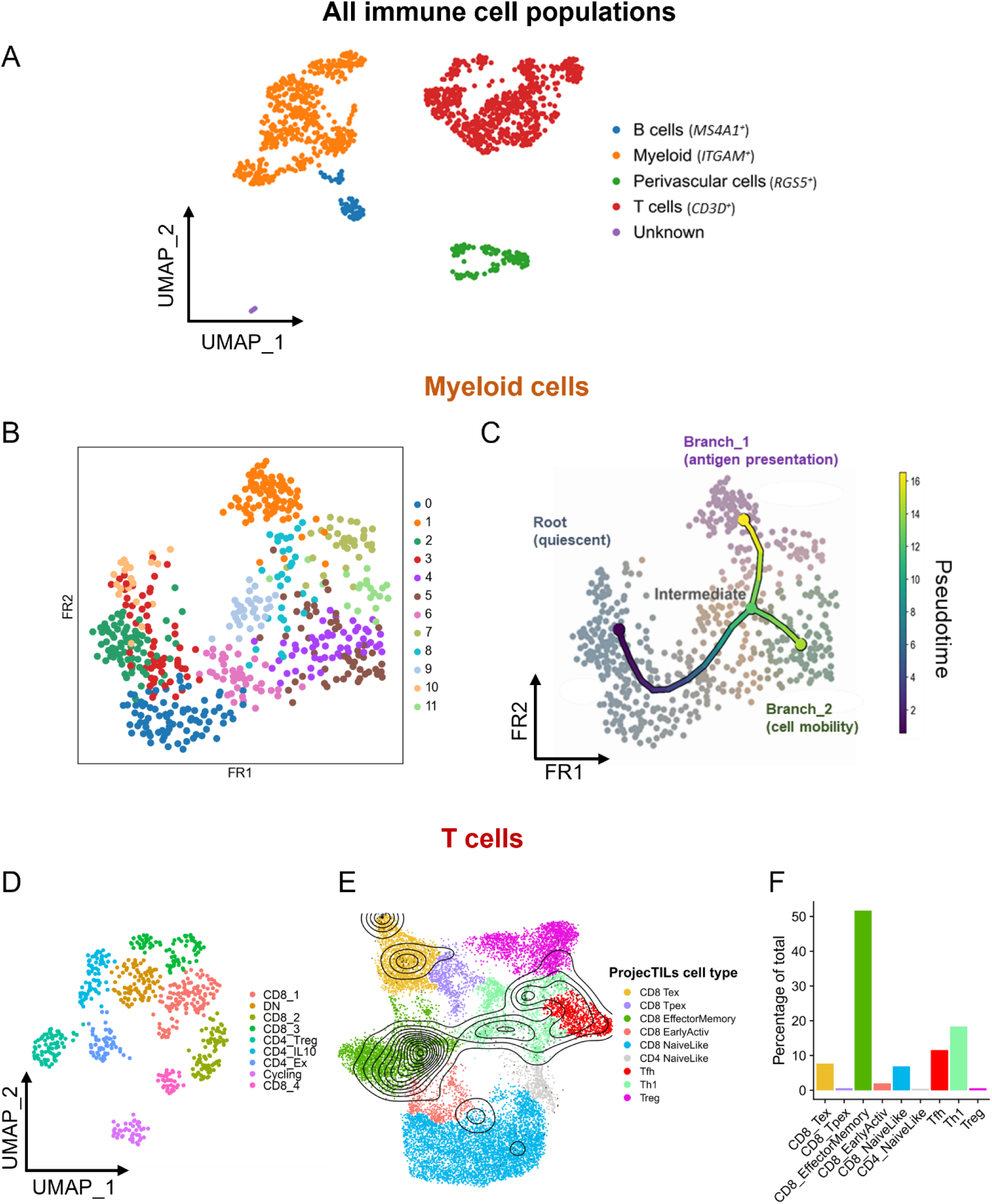
Single-cell characterisation of a type 1 cell-mediated response after AAV gene therapy in NHPs. (A) Identification of immune population cells in the PTPRC+ cluster. (B) Two-dimensional Fruchterman-Reingold (FR) force-directed graph of myeloid cells colored by unbiased Leiden cluster prior to trajectory inference analysis. (C) Pseudotime analysis of myeloid cells revealed one root (quiescent cells) and two branches – Branch 1 (antigen presentation) and Branch 2 (cell mobility). Gene ontology enrichment analysis (Figure S9) identified the main biological processes linked with these branches. (D) T cell (CD3D+) subclusters show presence of various CD4 and CD8 subsets, as well as a proliferating (‘cycling’) cluster with high expression of cell cycle genes. CD8+ subclusters labelled 1-4, DN – double negative (*CD4*-*CD8A*-). (E) Projection of our T cell dataset (black contour plot) onto the ProjecTILs reference dataset (colored UMAP). (F) Percentage of T cells projected onto each ProjecTILs cluster showing a prominent CD8 effector memory cell cluster.

To further characterise microglia/myeloid cell activation in response to AAV, we performed trajectory inference and pseudotime analysis on the *ITGAM*^+^ subcluster (**Figure 5 B-C** and **Figure S9**). The force-directed graph of myeloid cells showed a clear distinction between those present in all retinal areas and expressing homeostasis marker genes (*CX3CR1* and *P2RY12* high), and those present in the AAV8-GRK1-*mScarlet* and AAV8-GRK1-*hRPGRco* vector treated areas (**Figure S9 A-B**). Trajectory analysis suggested a bifurcation in myeloid cell trajectory over pseudotime whose termini we labelled as Branch 1 and Branch 2 (**Figure 5 B-C**). We did not find any of the clusters or either termini to be associated with a particular cell cycle phase, suggesting that cell division did not significantly influence the trajectory (**Figure S9 C**). We then performed statistical analysis of branch-specific genes and subsequent Gene Ontology (GO) enrichment analysis to identify branch-specific transcriptional programmes. We found that Branch 1 expressed a number of enriched GO gene sets largely associated with antigen presentation via both MHC class I and MHC class II, as well as viral response, lipid transport and other immune-related processes (**Figure S9 D-G**).

We next used a similar approach to identify subpopulations of T cells and found 4 clusters corresponding to CD8 T cells and 3 clusters corresponding to CD4 T cells (**Figure 5 D** and **Figure S10 A**). In addition, we found an apparent CD4-CD8-double negative (“DN”) cluster, as well as a proliferating (“cycling”) cluster with high S phase and G2M phase scores (**Figure S10 B**). To broadly understand the T cell states in AAV-treated retinas, we compared our data with ProjecTILs, a well-characterised and labelled T cell reference dataset, for classifying T cell subpopulations across species in the context of chronic viral infections ^31^ (**Figure 5 E**). The majority of CD8 T cells within our dataset (50%) projected onto the ‘CD8 Effector Memory’ cluster in ProjecTILs, with a smaller subset projecting onto the ‘CD8 Exhausted’ cluster. A small number of cells also projected onto the ‘CD8 Naïve-like’ cluster. The CD4 T cells mostly projected onto the Th1 and Tfh clusters, with a small number of Tregs (**Figure 5 F**). The CD8 T cells in AAV-treated retina showed high expression of cytotoxic genes (*GZMA*, *GZMB*, *GZMK* [Granzyme A, B, K], and *PRF1* [perforin]) with variation across clusters suggesting different activation states (**Figure S10 C**). One CD8 cluster also exhibited signs of exhaustion, with increased expression of *PRF1*, *CD8A* (cytotoxic T cell marker) and *TNFRSF9* (co-stimulatory receptor), as well as high expression of immune checkpoints: *PDCD1* (PD-1), *TIGIT*, *LAG3* and *HAVCR2* (TIM-3) (**Figure S10 D**). Among the CD4 T cells, we identified a Treg subset (*FOXP3, TIGIT, IKZF2*) and an *IL10*+ subset, possibly regulating AAV-induced inflammation. While *CXCR5* (indicative of T follicular helper cells) was low, some exhausted CD4 cells co-expressed *CXCL13, PDCD1* and *ICOS*, suggesting a small CXCR5lo T peripheral helper-like population (**Figure S10 D-F**).

Together, these findings suggest that subretinal AAV gene therapy in primates triggers a primarily type 1 cell-mediated response. By 3 months post-treatment, retinal CD8 effector memory T cells persist, alongside exhausted and naïve-like subsets. The concurrent presence of Th1 and Tfh-like CD4 populations indicates an antigen-experienced T cell milieu, potentially driving chronic retinal inflammation.

### Vitreous and retinal cytokine profile following subretinal gene therapy

Vitreous samples taken from each eye at baseline, 4, 8 and 12 weeks after subretinal gene therapy were analysed for inflammatory cytokines using the LEGENDplex NHP Inflammation Panel (**Figure 6 A and B**). This detected significant elevation of interferon gamma-induced protein-10 (IP-10 or CXCL10) in both NHPs and monocyte chemoattractant protein-1 (MCP-1 or CCL2) in NHP2. Both IP-10 and MCP-1 function as chemoattractant for monocytes/macrophages, T cells and NK cells, thus may contribute to persistent intraocular inflammation. All other cytokines in the NHP Inflammation Panel remained at minimal detectable levels at all time points, including TNFα, IFNβ, IFNγ, IL-23, IL-6, IL-8, IL-1β, IL10, IL-17a, IL12p40 and granulocyte-macrophage colony-stimulating factor (GM-CSF) (**Figure S11**). The notable absence of significant TNF-α response to AAV gene therapy may account for the lack of efficacy of adalimumab in the prevention of chronic GTAU. In addition, cytokine profiling of blood samples taken across the same timepoints did not show any significant change from baseline, indicating little systemic inflammatory effect from subretinal AAV gene therapy (**Figure S12**).

**Figure 6.**
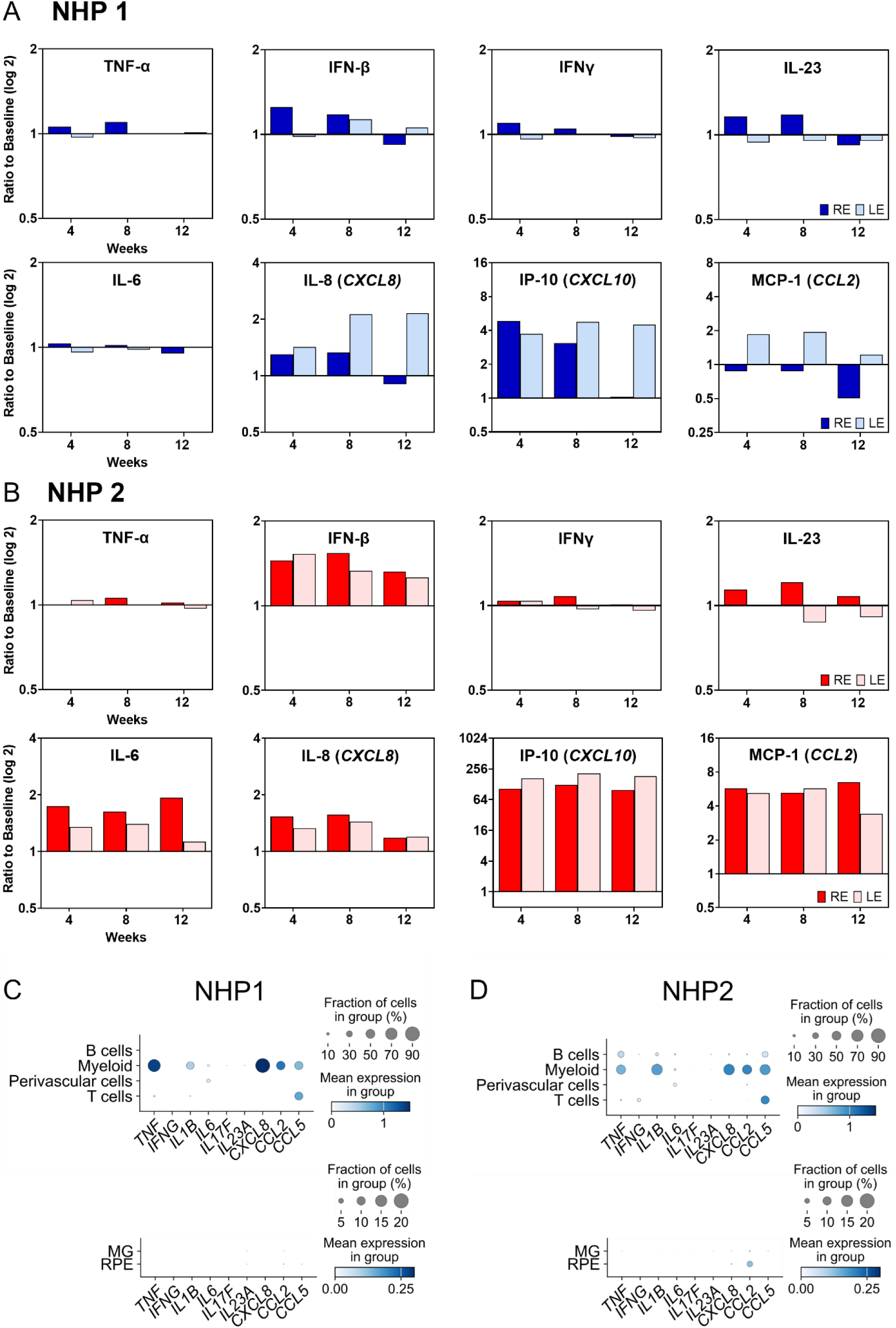
Cytokine expression profile after AAV gene therapy in NHPs. Quantification of vitreous cytokines in NHP1 (A) and NHP2 (B) using the LegendPlex NHP Inflammation Panel with levels expressed as fold-change (log 2) in mean fluorescence intensity (MFI) relative to baseline. RE = right eyes; LE = left eye. (C and D) Single-cell transcriptomic analysis of cytokine gene expression among retinal cell types in NHP1 and NHP2, respectively.

To identify which cells are responsible for the production of proinflammatory cytokines in GTAU, we further scrutinised the single-cell dataset. This revealed the myeloid cells in the retina as the dominant source of *MCP-1* (*CCL2*) expression as well as *IL-8 (or CXCL8,* a neutrophil chemoattractant), *CCL5* (a chemokine for T cells and macrophages) and *TNF α* (**Figure 6 C and D**). In contrast, the Müller glia and RPE cells, which may adopt active immune roles during retinal inflammation, did not demonstrate significant cytokine expression.

### iPSC-derived human microglia are rapidly activated by AAV *in vitro*

To validate whether microglia could be activated by AAV vectors, we conducted *in vitro* phagocytosis and cytokine assays using human iPSC-derived microglia (**Figure 7**). Phagocytic function was compared between (i) naïve microglia exposed to lipopolysaccharide (LPS) – positive control, (ii) naïve microglia exposed to AAV8, (iii) microglia pre-treated with AAV8 72 hours prior, and (iv) untreated naïve microglia – negative control. This showed rapid increases in phagocytosis in both LPS and AAV-treated microglia from 60 min onward, whereas previous exposure to AAV did not lead to persistent increase in phagocytic activity. Furthermore, we conducted cytokine profiling of the iPSC-microglia treated with AAV only versus AAV plus adalimumab (**Figure 7 C** and **Figure S13**). Microglia stimulated with AAV exhibited increased production of TNFα, IL-6, IL-8 and MCP-1 between 1 to 72 hours. Adalimumab treatment mitigated the TNFα spike but had little to no effect on the other cytokines, thus indicating a selective but limited modulatory effect on AAV-induced inflammation. Interestingly, MCP-1 was once again detected at high levels in both the AAV only and AAV plus adalimumab arms, consistent with activated microglia being a major producer of MCP-1. Taken together, the results suggest that microglia possess pattern recognition receptors that can detect the presence of AAV particles and activate a rapid innate immune response that contributes to the recruitment of adaptive immunity.

**Figure 7.**
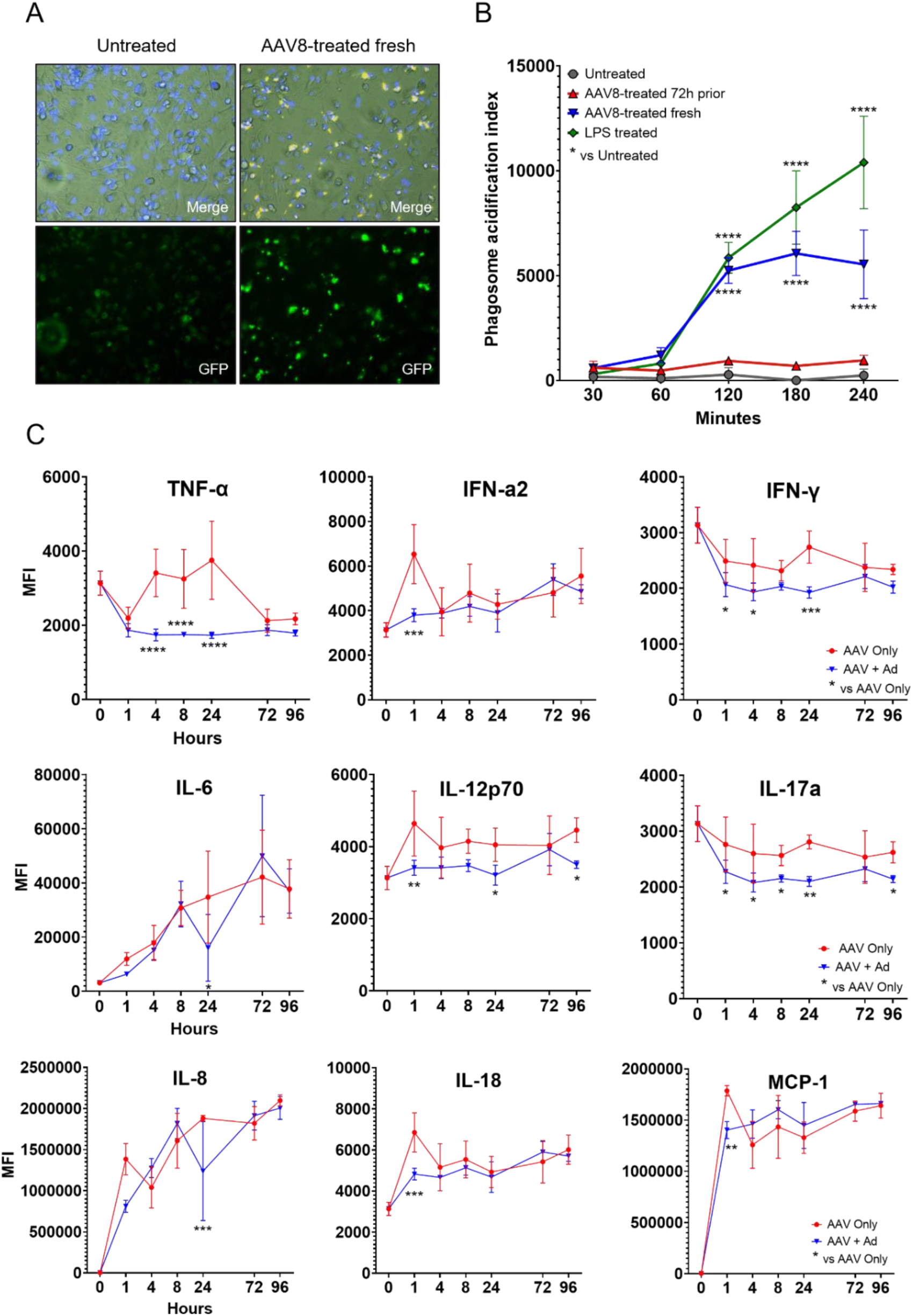
Activation of human iPSC-derived microglia by AAV exposure. (A) Representative fluorescence microscopy images showing phagocytosis of pHrodo-labelled zymosan particles by iPSC-microglia exposed to AAV at 240 min. pHrodo becomes fluorescent upon acidification within endosomes. (B) Phagocytic assay comparing (i) naïve microglia exposed to lipopolysaccharide (LPS) (green), (ii) naïve microglia exposed to AAV8-CAG-mScarlet (blue), (iii) microglia pre-treated with AAV8 72 hours prior (red), and (iv) untreated naïve microglia (grey). The phagosome acidification index was calculated as the total amount of fluorescence signal above threshold over the number of cells. (C) Cytokine profile of iPSC-microglia treated with AAV only versus AAV + adalimumab. Errors bars represent SD (n=4). Two-way ANOVA test was performed: * p>0.01; ** 0.001<p<0.01; *** 0.0001<p<0.001; **** p<0.0001.

## DISCUSSION

In this study, we simulated subretinal gene therapy in primates using human therapeutic AAV vector doses and surgical technique, and conducted deep assessment of the resulting retinal cellular responses at single cell resolution. Using single-cell sequencing and spatial transcriptomic approaches, we were able to distinguish between viral transgene expression and native gene expression in wildtype animals and identify subpopulations of immune cells to interrogate the key immune interactions involved in gene therapy-associated uveitis (GTAU).

The results demonstrate that both clinical-grade AAV2-CAG-*hRPE65* (Luxturna) and engineering-grade AAV8-GRK1-*hRPGRco* were well tolerated *in vivo* with no adverse effects on retinal structure and function. This aligns with the safety profiles of both vectors reported in clinical trials ^2,4^. Within the subretinal bleb area, the AAV2 vector containing the ubiquitous CAG promoter transduced approximately 30% of RPE – the target cell population for functional rescue in *RPE65*-associated Leber congenital amaurosis (LCA). This level of transduction could potentially impose a limit on visual improvement as 70% of RPE cells remain deficient for *RPE65* thus unable to complete the visual cycle. In addition, the single-cell data shows 30% transduction of rod photoreceptors. While such ectopic expression of *RPE65* may be unhelpful in treating LCA, it could be a desirable characteristic for other ‘biofactory’ gene augmentation approaches aimed at producing a secreted protein (e.g. AAV2-CAG-*complement factor I* gene therapy for dry AMD) ^40^. By contrast, the AAV8 vector containing the photoreceptor-specific GRK1 promoter transduced 70-80% of rods and cones, which represents the majority of the target cell types in *RPGR*-associated X-linked retinitis pigmentosa (XLRP). This could account for the dramatic microperimetry retinal sensitivity improvements seen in clinical trials ^3,4,41^. In this instance, the GRK1 promoter robustly restricted transgene expression to photoreceptors, thus minimising off-target effects. Moreover, spatial transcriptomic data indicate the level of AAV-mediate human *RPGR* transgene expression was considerably greater than native macaque *RPGR* expression (**Figure 3 E)**. Overexpression of RPGR has previously been reported to be associated with protein mislocalization and potential retinal toxicity in preclinical studies ^42–44^. However, we did not find any correlation between the level of transgene expression and upregulation of apoptotic marker genes (*BAX*, *CASP3* and *APAF1*) at the single cell level. This somewhat abrogates concerns that transgene overexpression may be a cause of retinal toxicity at the therapeutic dose ^45^.

A number of clinical studies have reported incidence of chorioretinal atrophy (CRA) following retinal gene therapy with voretigene neparvovec (Weed et al. 2019; Gange et al. 2022; Fischer et al. 2024; Kvanta et al. 2024). The mechanism of CRA remains unclear. In NHP2, we noted CRA development (seen as confluent hypo-autofluorescence) around the area treated with laboratory-grade AAV8-GRK1-*mScarlet* reporter vector (**Figure 2 B** left eye), which was associated with subretinal infiltrates on OCT and the greatest contribution of immune cells to the single-cell analysis (**Figure S8 C**). This suggest that retinal inflammation can be a potential cause of chorioretinal atrophy. Single-cell analysis of immune cells from the *mScarlet* vector-treated area revealed upregulation of MHC class-I and immunoproteosome-related genes in rods and cones (suggesting increased presentation of intracellular antigens) as well as anti-viral response genes (**Figure S7**). For instance, this could promote presentation of capsid and transgene peptides to infiltrating CD8 T cells, triggering targeted photoreceptor destruction. We also observed a Type I interferon response with upregulation of IFNα/β receptor subunit (*IFNAR2*), interferon regulatory factor (*IRF7*), interferon-related genes (*DDX60, OAS1, OAS3*) in rods, as well as elevated IFNβ in the vitreous (**Figure 6**). This could lead to suppression of transgene expression and stimulate myeloid/NK cell anti-viral responses. Interestingly, while immune cell infiltration and CRA were limited to the inferior retina treated with the AAV8-GRK1-*mScarlet* vector, the superior retina treated with a similar AAV8-GRK1-*RPGR* vector and untreated nasal retina remained quiescent with little evidence of cross-reactivity or generalised blood-retinal barrier breakdown. Increased immunogenicity of mScarlet may be due to its exogenous nature, whereas human and macaque RPGR show high level of amino acid sequence homology except for indels within the highly repetitive C-terminus encoded by ORF15 ^46^. Alternatively, laboratory-grade vectors may contain impurities such as residual host cell proteins or DNA fragments, which could play a role in the activation of innate immunity.

Our primate transcriptomic data suggest that the retinal immune infiltrate in GTAU is predominantly a type I cell-mediated response consisting of myeloid cells and T cells (**Figure 5** and **S6 B, D**). This is consistent with previous observations in mice, where subretinal injection of AAV8-CAG-*GFP* led to a similar composition of leukocyte infiltrate detected by flow cytometry ^15^. Comparison between single-cell data from untreated and treated retinal punches strongly implicates activation of resident microglia by AAV vectors (**Figure 5** and **S8**). Trajectory analysis infers transition of myeloid cells (including microglia) from a resting state towards two activated states characterised by upregulation of (i) antigen presentation and (ii) cell motility gene sets. An acquired antigen presenting function in microglia would be expected to restimulate AAV antigen-specific infiltrating T cells. Enhanced mobility would enable migration of microglia to the subretinal space, a phenomenon previously suggested by Xiong et al. 2019 ^47^ and corroborated by our spatial transcriptomic mapping (**Figure 4**). Whether the two branches based on trajectory inference are truly separate, or whether the motility gene expression programme is an intermediate on the path to an antigen presenting phenotype requires further exploration. Further insights into the role of microglia in GTAU were gained from human iPSC-derived microglia stimulated with AAV vectors. AAV exposure led to a rapid increase in microglia phagocytic activity and release of inflammatory cytokines, including TNFα, IFNα2, IL-6 and MCP-1. MCP-1 was consistently identified as a key monocyte chemoattractant expressed by myeloid cells in the NHP retinas by single cell analysis and a prominent cytokine in vitreous biopsies. Together, these results implicate retinal microglia as a key orchestrator of immune response against AAV and recruiter of adaptive immunity.

At the 12-week timepoint post-gene therapy, the infiltrating T cells in the retina showed signs of exhaustion similar to those seen during chronic viral infection, with high expression of cytotoxicity genes and immune checkpoints (*PD1*, *TIGIT* and *ICOS*) (**Figure S10 D** and **E**). We also detected a subset of CD8+ effector memory T cells. The extent to which these cells are actively killing transduced retinal cells remain to be determined. Their presence carries significant clinical implications for second eye treatment in the same patient as re-challenge with the same antigen may lead to more severe and prolonged retinal inflammation. Interestingly, we also found evidence of both Tregs and IL10+ T cells, which are likely dampening the ongoing immune response. Understanding the interactions between these subsets of T cells and the myeloid cells will be important in developing more effective prophylaxis and treatment for GTAU. The use of adjunctive intravitreal adalimumab at 4 weekly intervals had limited impact on the immune response seen *in vivo* at 12 weeks over and above the effects of a single dose of periocular corticosteroid (triamcinolone) at the time of gene therapy surgery. While adalimumab did suppress the transient TNFα rise (over 72 hours) in iPSC-microglia exposed to AAV *in vitro* (**Figure 7**), the paucity of TNFα in vitreous samples from both NHPs suggests that it may not be a major cytokine driving chronic GTAU (**Figure 6**). Future studies may benefit from targeting alternative immune signalling pathways specific to GTAU.

In conclusion, our appreciation of the cellular effects of AAV gene therapy in human eyes has so far been limited by the ‘black box’ effect of the treatment in patients. This study provides deep insight into the nature of immune response to AAV-mediated retinal gene therapy in primates using a combination of spatial and single-cell transcriptomic approaches. Spatial transcriptomics has gained significant traction in recent years to bridge the gap between structural and transcriptomic analyses, but has seen limited application to the retina to date in the mouse ^48^ and human ^49^. To our knowledge, the present study is the first to apply spatial transcriptomics to the NHP retina. These large datasets encompassing both treated and untreated primate retinas provide an atlas for identifying key immune interactions that may be targeted to mitigate GTAU.

## AUTHOR CONTRIBUTIONS

KX conceptualized, planned and supervised the project. CMFC provided AAV vectors and performed tissue processing for spatial analysis. KX performed gene therapy surgery with assistance from JCK and REM. CS and SC produced iPSC-derived microglia. MJ produced the *mScarlet* vector and conducted immunological analyses. JQ performed single-cell RNA-sequencing and bioinformatic analysis. CS performed immunohistochemistry, spatial transcriptomic processing and analysis. CS and KX wrote the original manuscript, and all authors contributed to the editing and reviewing process.

## ACKNOWLEDGEMENTS

This work was supported by grant funding from the Wellcome Trust (216593/Z/19/Z). JQ and JCK would like to acknowledge funding from the Medical Research Council (MRC). MCJ would like to acknowledge funding from the Biotechnology & Biological Sciences Research Council (BBSRC). KX is indebted to Prof Alistair Lamb (Nuffield Department of Surgical Sciences, University of Oxford) for advice on spatial transcriptomic analysis, Prof Andrew Dick (UCL Institute of Ophthalmology) and Prof M Dominik Fischer (Nuffield Department of Clinical Neurosciences, University of Oxford) for discussions over NHP experimental plans. We thank the staff members of Biomedical Services, Lucy Underdown, Dr Henri Bertrand, Katie Underdown, Sarah Rohling, Andrew Emberton and Kelly Simpson for their expert care of the animals and assistance with anaesthesia. We are also grateful to Dr Helen Ferry at the Flow Sorting Facility (Experimental Medicine Division, University of Oxford) for expert technical assistance with flow cytometric analysis.

## REFERENCES

1. Purdy, R., John, M., Bray, A., Clare, A.J., Copland, D.A., Chan, Y.K., Henderson, R.H., Nerinckx, F., Leroy, B.P., Yang, P., Pennesi, M.E., et al. (2025). Gene Therapy-Associated Uveitis (GTAU): Understanding and mitigating the adverse immune response in retinal gene therapy. Progress in Retinal and Eye Research 106, 101354. 10.1016/j.preteyeres.2025.101354.

2. Russell, S., Bennett, J., Wellman, J.A., Chung, D.C., Yu, Z.F., Tillman, A., Wittes, J., Pappas, J., Elci, O., McCague, S., Cross, D., et al. (2017). Efficacy and safety of voretigene neparvovec (AAV2-hRPE65v2) in patients with RPE65-mediated inherited retinal dystrophy: a randomised, controlled, open-label, phase 3 trial. Lancet 390, 849–860. 10.1016/s0140-6736(17)31868-8.

3. Cehajic-Kapetanovic, J., Xue, K., Martinez-Fernandez de la Camara, C., Nanda, A., Davies, A., Wood, L.J., Salvetti, A.P., Fischer, M.D., Aylward, J.W., Barnard, A.R., Jolly, J.K., et al. (2020). Initial results from a first-in-human gene therapy trial on X-linked retinitis pigmentosa caused by mutations in RPGR. Nat Med 26, 354–359. 10.1038/s41591-020-0763-1.

4. Lam, B.L., Pennesi, M.E., Kay, C.N., Panda, S., Gow, J.A., Zhao, G., and MacLaren, R.E. (2024). Assessment of Visual Function with Cotoretigene Toliparvovec in X-Linked Retinitis Pigmentosa in the Randomized XIRIUS Phase 2/3 Study. Ophthalmology 131, 1083–1093. 10.1016/j.ophtha.2024.02.023.

5. Yang, P., Birch, D., Lauer, A., Sisk, R., Anand, R., Pennesi, M.E., Iannaccone, A., Yaghy, A., Scaria, A., Jung, J.A., Curtiss, D., et al. (2025). Subretinal Gene Therapy Drug AGTC-501 for XLRP Phase 1/2 Multicenter Study (HORIZON): 24-Month Safety and Efficacy Results. Am J Ophthalmol 271, 268–285. 10.1016/j.ajo.2024.11.021.

6. Fischer, M.D., Ochakovski, G.A., Beier, B., Seitz, I.P., Vaheb, Y., Kortuem, C., Reichel, F.F.L., Kuehlewein, L., Kahle, N.A., Peters, T., Girach, A., et al. (2019). Efficacy and Safety of Retinal Gene Therapy Using Adeno-Associated Virus Vector for Patients With Choroideremia: A Randomized Clinical Trial. JAMA Ophthalmology 137, 1247–1254. 10.1001/jamaophthalmol.2019.3278.

7. Fischer, M.D., Simonelli, F., Sahni, J., Holz, F.G., Maier, R., Fasser, C., Suhner, A., Stiehl, D.P., Chen, B., Audo, I., and Leroy, B.P. (2024). Real-World Safety and Effectiveness of Voretigene Neparvovec: Results up to 2 Years from the Prospective, Registry-Based PERCEIVE Study. Biomolecules.

8. MacLaren, R.E., Fischer, M.D., Gow, J.A., Lam, B.L., Sankila, E.-M.K., Girach, A., Panda, S., Yoon, D., Zhao, G., and Pennesi, M.E. (2023). Subretinal timrepigene emparvovec in adult men with choroideremia: a randomized phase 3 trial. Nature Medicine 29, 2464–2472. 10.1038/s41591-023-02520-3.

9. Pierce Eric, A., Aleman Tomas, S., Jayasundera Kanishka, T., Ashimatey Bright, S., Kim, K., Rashid, A., Jaskolka Michael, C., Myers Rene, L., Lam Byron, L., Bailey Steven, T., Comander Jason, I., et al. (2024). Gene Editing for CEP290-Associated Retinal Degeneration. New England Journal of Medicine 390, 1972–1984. 10.1056/NEJMoa2309915.

10. Campochiaro, P.A., Avery, R., Brown, D.M., Heier, J.S., Ho, A.C., Huddleston, S.M., Jaffe, G.J., Khanani, A.M., Pakola, S., Pieramici, D.J., Wykoff, C.C., et al. (2024). Gene therapy for neovascular age-related macular degeneration by subretinal delivery of RGX-314: a phase 1/2a dose-escalation study. The Lancet 403, 1563–1573. 10.1016/S0140-6736(24)00310-6.

11. Kessel, L., Christensen, U.C., and Klemp, K. (2022). Inflammation after Voretigene Neparvovec Administration in Patients with RPE65-Related Retinal Dystrophy. Ophthalmology 129, 1287–1293. 10.1016/j.ophtha.2022.06.018.

12. Yang, P., Pardon, L.P., Ho, A.C., Lauer, A.K., Yoon, D., Boye, S.E., Boye, S.L., Roman, A.J., Wu, V., Garafalo, A.V., Sumaroka, A., et al. (2024). Safety and efficacy of ATSN-101 in patients with Leber congenital amaurosis caused by biallelic mutations in GUCY2D: a phase 1/2, multicentre, open-label, unilateral dose escalation study. The Lancet 404, 962–970. 10.1016/S0140-6736(24)01447-8.

13. Reichel, F.F., Seitz, I., Wozar, F., Dimopoulos, S., Jung, R., Kempf, M., Kohl, S., Kortüm, F.C., Ott, S., Pohl, L., Stingl, K., et al. (2023). Development of retinal atrophy after subretinal gene therapy with voretigene neparvovec. British Journal of Ophthalmology 107, 1331. 10.1136/bjophthalmol-2021-321023.

14. Seitz, I.P., Wozar, F., Ochakovski, G.A., Reichel, F.F., Gelisken, F., Bartz-Schmidt, K.U., Peters, T., and Fischer, M.D. (2024). Dose-Dependent Progression of Chorioretinal Atrophy at the Injection Site After Subretinal Injection of rAAV2/8 in Nonhuman Primates. Ophthalmology Science 4, 100516. 10.1016/j.xops.2024.100516.

15. Chandler, L.C., McClements, M.E., Yusuf, I.H., Martinez-Fernandez de la Camara, C., MacLaren, R.E., and Xue, K. (2021). Characterizing the cellular immune response to subretinal AAV gene therapy in the murine retina. Mol Ther Methods Clin Dev 22, 52–65. 10.1016/j.omtm.2021.05.011.

16. Durrani, K., Kempen, J.H., Ying, G.S., Kacmaz, R.O., Artornsombudh, P., Rosenbaum, J.T., Suhler, E.B., Thorne, J.E., Jabs, D.A., Levy-Clarke, G.A., Nussenblatt, R.B., et al. (2017). Adalimumab for Ocular Inflammation. Ocul Immunol Inflamm 25, 405–412. 10.3109/09273948.2015.1134581.

17. Bober, E., Frain, K., Fotuhi, M., Virgo, J., Hindle, E., Ma, J., Luis, J., Addison, P., Okhravi, N., Tucker, W., Westcott, M., et al. (2024). Adalimumab in the treatment of refractory non-infectious scleritis: 6-month outcomes. Eye 38, 628–630. 10.1038/s41433-023-02725-3.

18. Çam, F., and Celiker, H. (2024). Efficacy, retention rate and safety of adalimumab treatment in patients with non-infectious uveitis and scleritis: a real-world, retrospective, single-centre study. Eye 38, 893–901. 10.1038/s41433-023-02800-9.

19. Xue, K., Groppe, M., Salvetti, A.P., and MacLaren, R.E. (2017). Technique of retinal gene therapy: delivery of viral vector into the subretinal space. Eye (Lond) 31, 1308–1316. 10.1038/eye.2017.158.

20. Snodderly, D.M., Sandstrom, M.M., Leung, I.Y.F., Zucker, C.L., and Neuringer, M. (2002). Retinal Pigment Epithelial Cell Distribution in Central Retina of Rhesus Monkeys. Investigative Ophthalmology & Visual Science 43, 2815–2818.

21. Schindelin, J., Arganda-Carreras, I., Frise, E., Kaynig, V., Longair, M., Pietzsch, T., Preibisch, S., Rueden, C., Saalfeld, S., Schmid, B., Tinevez, J.-Y., et al. (2012). Fiji: an open-source platform for biological-image analysis. Nature Methods 9, 676–682. 10.1038/nmeth.2019.

22. Young, M.D., and Behjati, S. (2020). SoupX removes ambient RNA contamination from droplet-based single-cell RNA sequencing data. Gigascience 9. 10.1093/gigascience/giaa151.

23. Germain, P.L., Lun, A., Garcia Meixide, C., Macnair, W., and Robinson, M.D. (2021). Doublet identification in single-cell sequencing data using scDblFinder. F1000Res 10, 979. 10.12688/f1000research.73600.2.

24. Wolf, F.A., Angerer, P., and Theis, F.J. (2018). SCANPY: large-scale single-cell gene expression data analysis. Genome Biol 19, 15. 10.1186/s13059-017-1382-0.

25. Hao, Y., Stuart, T., Kowalski, M.H., Choudhary, S., Hoffman, P., Hartman, A., Srivastava, A., Molla, G., Madad, S., Fernandez-Granda, C., and Satija, R. (2024). Dictionary learning for integrative, multimodal and scalable single-cell analysis. Nature Biotechnology 42, 293–304. 10.1038/s41587-023-01767-y.

26. Badia, I.M.P., Vélez Santiago, J., Braunger, J., Geiss, C., Dimitrov, D., Müller-Dott, S., Taus, P., Dugourd, A., Holland, C.H., Ramirez Flores, R.O., and Saez-Rodriguez, J. (2022). decoupleR: ensemble of computational methods to infer biological activities from omics data. Bioinform Adv 2, vbac016. 10.1093/bioadv/vbac016.

27. Love, M.I., Huber, W., and Anders, S. (2014). Moderated estimation of fold change and dispersion for RNA-seq data with DESeq2. Genome Biol 15, 550. 10.1186/s13059-014-0550-8.

28. Sergushichev, A.A. (2016). An algorithm for fast preranked gene set enrichment analysis using cumulative statistic calculation. bioRxiv, 060012. 10.1101/060012.

29. Faure, L., Soldatov, R., Kharchenko, P.V., and Adameyko, I. (2023). scFates: a scalable python package for advanced pseudotime and bifurcation analysis from single-cell data. Bioinformatics 39. 10.1093/bioinformatics/btac746.

30. Fruchterman, T.M.J., and Reingold, E.M. (1991). Graph drawing by force-directed placement. Software: Practice and Experience 21, 1129–1164. 10.1002/spe.4380211102.

31. Andreatta, M., Corria-Osorio, J., Müller, S., Cubas, R., Coukos, G., and Carmona, S.J. (2021). Interpretation of T cell states from single-cell transcriptomics data using reference atlases. Nat Commun 12, 2965. 10.1038/s41467-021-23324-4.

32. Washer, S.J., Perez-Alcantara, M., Chen, Y., Steer, J., James, W.S., Trynka, G., Bassett, A.R., and Cowley, S.A. (2022). Single-cell transcriptomics defines an improved, validated monoculture protocol for differentiation of human iPSC to microglia. Sci Rep 12, 19454. 10.1038/s41598-022-23477-2.

33. Suhler, E.B., Adán, A., Brézin, A.P., Fortin, E., Goto, H., Jaffe, G.J., Kaburaki, T., Kramer, M., Lim, L.L., Muccioli, C., Nguyen, Q.D., et al. (2018). Safety and Efficacy of Adalimumab in Patients with Noninfectious Uveitis in an Ongoing Open-Label Study: VISUAL III. Ophthalmology 125, 1075–1087. 10.1016/j.ophtha.2017.12.039.

34. Brouwer, A.H., de Wit, G.C., de Boer, J.H., and van Genderen, M.M. (2020). Effects of DTL electrode position on the amplitude and implicit time of the electroretinogram. Doc Ophthalmol 140, 201–209. 10.1007/s10633-019-09733-3.

35. Brouwer, A.H., de Wit, G.C., Ten Dam, N.H., Wijnhoven, R., van Genderen, M.M., and de Boer, J.H. (2019). Prolonged Cone b-Wave on Electroretinography Is Associated with Severity of Inflammation in Noninfectious Uveitis. Am J Ophthalmol 207, 121–129. 10.1016/j.ajo.2019.05.028.

36. Berson, E.L., Sandberg, M.A., Rosner, B., Birch, D.G., and Hanson, A.H. (1985). Natural course of retinitis pigmentosa over a three-year interval. Am J Ophthalmol 99, 240–251. 10.1016/0002-9394(85)90351-4.

37. Beltran, W.A., Cideciyan, A.V., Boye, S.E., Ye, G.-J., Iwabe, S., Dufour, V.L., Marinho, L.F., Swider, M., Kosyk, M.S., Sha, J., Boye, S.L., et al. (2017). Optimization of Retinal Gene Therapy for X-Linked Retinitis Pigmentosa Due to RPGR Mutations. Molecular Therapy 25, 1866–1880. 10.1016/j.ymthe.2017.05.004.

38. Quinn, J., Salman, A., Paluch, C., Jackson-Wood, M., McClements, M.E., Luo, J., Davis, S.J., Cornall, R.J., MacLaren, R.E., Dendrou, C.A., and Xue, K. (2024). Single-cell transcriptomic analysis of retinal immune regulation and blood-retinal barrier function during experimental autoimmune uveitis. Sci Rep 14, 20033. 10.1038/s41598-024-68401-y.

39. Wiley, L.A., Boyce, T.M., Meyering, E.E., Ochoa, D., Sheehan, K.M., Stone, E.M., Mullins, R.F., Tucker, B.A., and Han, I.C. (2023). The Degree of Adeno-Associated Virus-Induced Retinal Inflammation Varies Based on Serotype and Route of Delivery: Intravitreal, Subretinal, or Suprachoroidal. Human Gene Therapy 34, 530–539. 10.1089/hum.2022.222.

40. Hallam, T.M., Gardenal, E., McBlane, F., Cho, G., Ferraro, L.L., Pekle, E., Lu, D., Carney, K., Wenden, C., Beadsmoore, H., Kaiser, S., et al. (2025). Ocular biomarker profiling after complement factor I gene therapy in geographic atrophy secondary to age-related macular degeneration. 10.7554/elife.99806.2.

41. von Krusenstiern, L., Liu, J., Liao, E., Gow, J.A., Chen, G., Ong, T., Lotery, A.J., Jalil, A., Lam, B.L., and MacLaren, R.E. (2023). Changes in Retinal Sensitivity Associated With Cotoretigene Toliparvovec in X-Linked Retinitis Pigmentosa With RPGR Gene Variations. JAMA Ophthalmol 141, 275–283. 10.1001/jamaophthalmol.2022.6254.

42. Wright, R.N., Hong, D.H., and Perkins, B. (2011). Misexpression of the constitutive Rpgr(ex1-19) variant leads to severe photoreceptor degeneration. Invest Ophthalmol Vis Sci 52, 5189–5201. 10.1167/iovs.11-7470.

43. Dufour, V.L., Cideciyan, A.V., Ye, G.J., Song, C., Timmers, A., Habecker, P.L., Pan, W., Weinstein, N.M., Swider, M., Durham, A.C., Ying, G.S., et al. (2020). Toxicity and Efficacy Evaluation of an Adeno-Associated Virus Vector Expressing Codon-Optimized RPGR Delivered by Subretinal Injection in a Canine Model of X-linked Retinitis Pigmentosa. Hum Gene Ther 31, 253–267. 10.1089/hum.2019.297.

44. Song, C., Dufour, V.L., Cideciyan, A.V., Ye, G.J., Swider, M., Newmark, J.A., Timmers, A.M., Robinson, P.M., Knop, D.R., Chulay, J.D., Jacobson, S.G., et al. (2020). Dose Range Finding Studies with Two RPGR Transgenes in a Canine Model of X-Linked Retinitis Pigmentosa Treated with Subretinal Gene Therapy. Hum Gene Ther 31, 743–755. 10.1089/hum.2019.337.

45. Stingl, K., Stingl, K., Schwartz, H., Reid, M.W., Kempf, M., Dimopoulos, S., Kortuem, F., Borchert, M.S., Lee, T.C., and Nagiel, A. (2023). Full-field Scotopic Threshold Improvement after Voretigene Neparvovec-rzyl Treatment Correlates with Chorioretinal Atrophy. Ophthalmology 130, 764–770. 10.1016/j.ophtha.2023.02.015.

46. Raghupathy, R.K., Gautier, P., Soares, D.C., Wright, A.F., and Shu, X. (2015). Evolutionary Characterization of the Retinitis Pigmentosa GTPase Regulator Gene. Invest Ophthalmol Vis Sci 56, 6255–6264. 10.1167/iovs.15-17726.

47. Xiong, W., Wu, D.M., Xue, Y., Wang, S.K., Chung, M.J., Ji, X., Rana, P., Zhao, S.R., Mai, S., and Cepko, C.L. (2019). AAV cis-regulatory sequences are correlated with ocular toxicity. Proc Natl Acad Sci U S A 116, 5785–5794. 10.1073/pnas.1821000116.

48. Choi, J., Li, J., Ferdous, S., Liang, Q., Moffitt, J.R., and Chen, R. (2023). Spatial organization of the mouse retina at single cell resolution by MERFISH. Nature Communications 14, 4929. 10.1038/s41467-023-40674-3.

49. Zhang, J., Wang, J., Zhou, Q., Chen, Z., Zhuang, J., Zhao, X., Gan, Z., Wang, Y., Wang, C., Molday, R.S., Yang, Y.T., et al. (2025). Spatiotemporally resolved transcriptomics reveals the cellular dynamics of human retinal development. Nature Communications 16, 2307. 10.1038/s41467-025-57625-9.

