## Supplemental Material (Figures S1 to S12, Tables S1 and S2) for "Single-cell and spatial transcriptomic analyses of gene therapy-associated retinal inflammation in non-human primates"

***Figure S1: No deterioration of the macula after AAV gene therapy in NHPs.***
See Figure 2. Longitudinal multimodal retinal imaging of the macula following subretinal injection of AAV vectors over 12 weeks of both eyes NHP1 (**A**) and NHP2 (**B**). The colour lines represent the locations of OCT sections from inside the treated subretinal blebs. Note that all retinal images are vertically inverted (top of the image representing inferior retina). Scale bars = 500 µm. (**C**) Retina thickness surrounding the macula extracted from the OCT images.

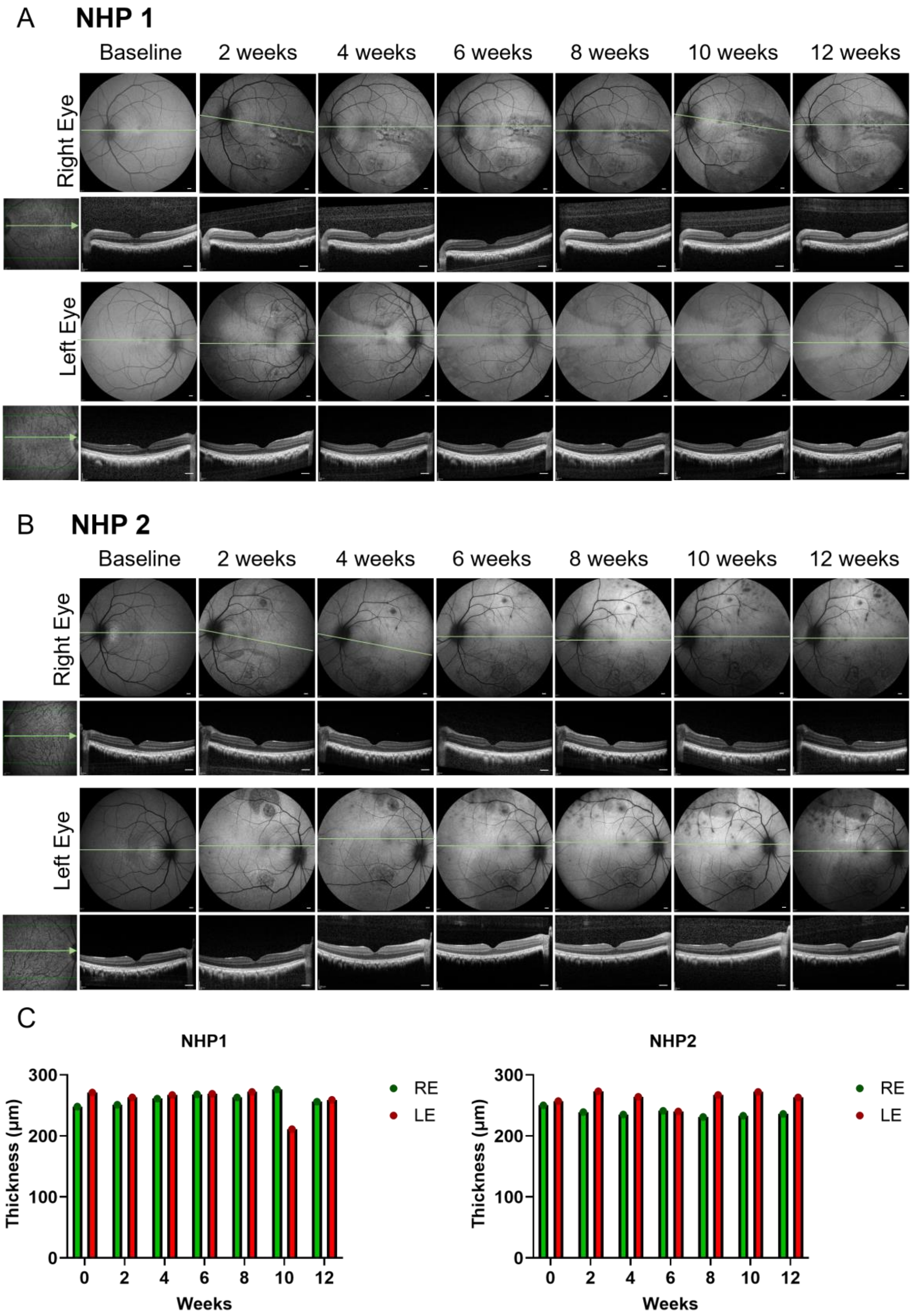

***Figure S2: NHP1 ERG data.***

Electroretinography (ERG) responses recorded under dark-adapted (DA) and light-adapted (LA) conditions at baseline and after 10 weeks for RE and LE of NHP1. Measurements include standard a-wave and b-wave responses, as well as oscillatory potentials (OPs) and 30 Hz flicker ERG to assess retinal function. Performed in duplicates.

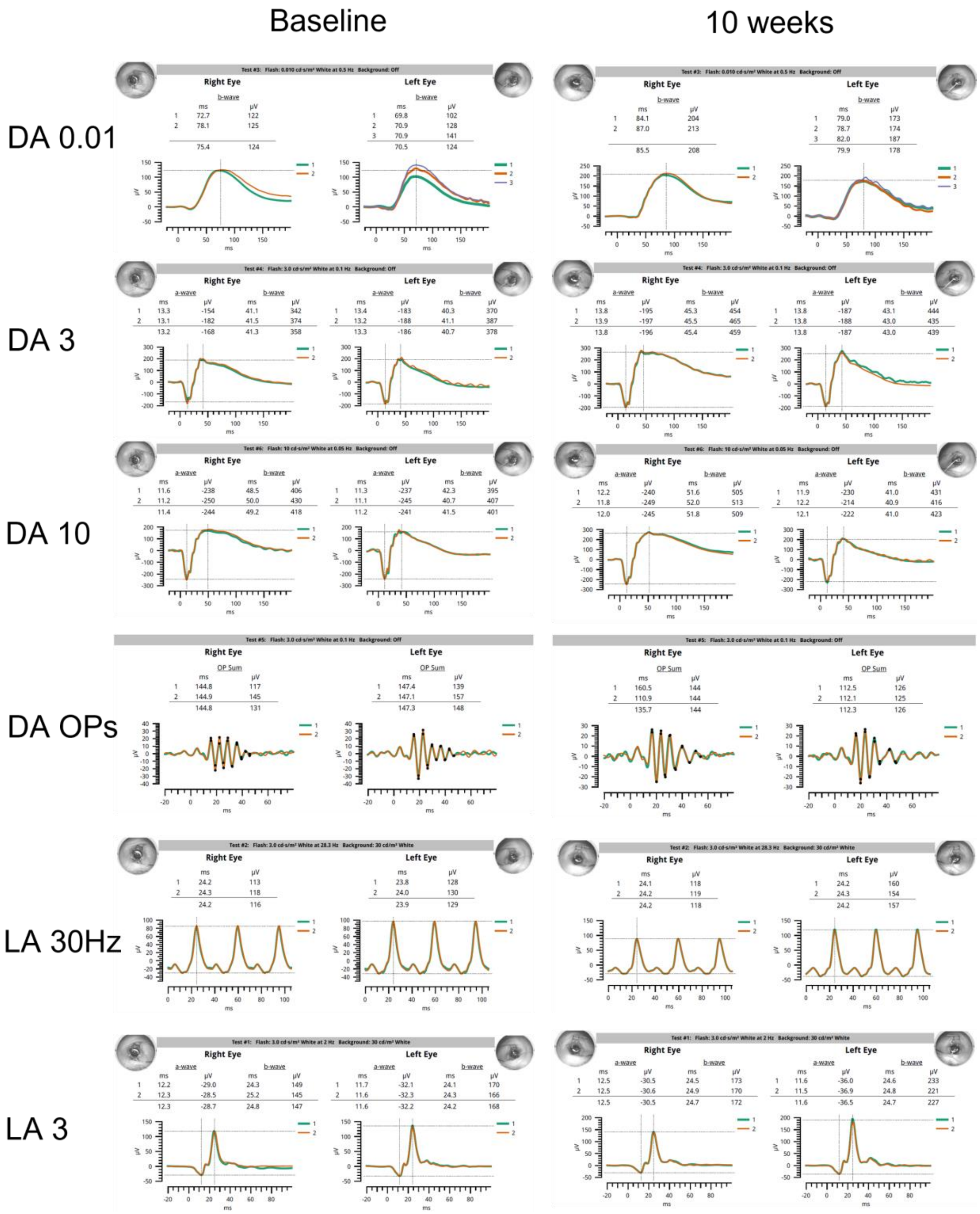

***Figure S3: NHP2 ERG data.***

Electroretinography (ERG) responses recorded under dark-adapted (DA) and light-adapted (LA) conditions at baseline and after 10 weeks for RE and LE of NHP2. Measurements include standard a-wave and b-wave responses, as well as oscillatory potentials (OPs) and 30 Hz flicker ERG to assess retinal function. Performed in duplicates.

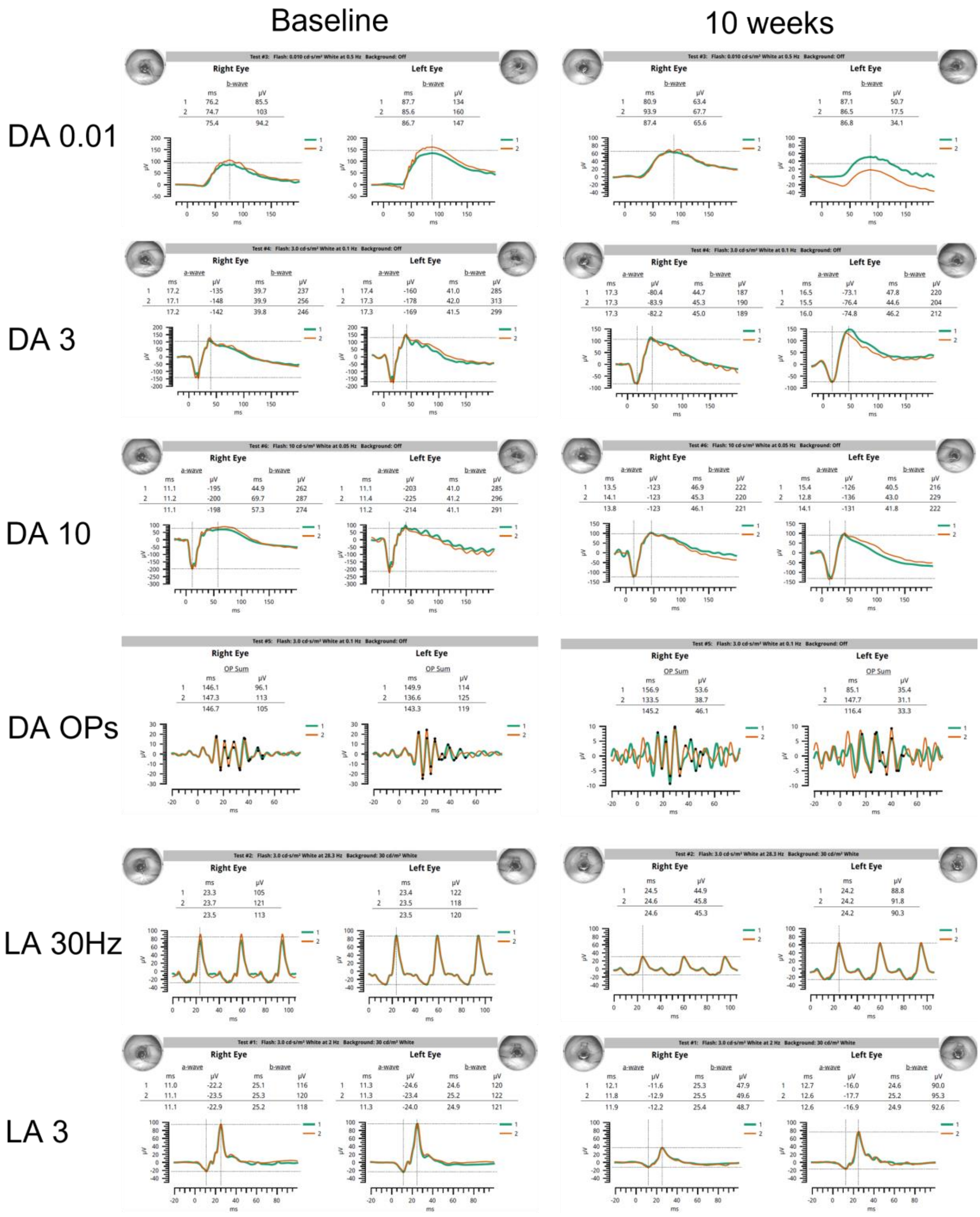

***Figure S4: Single-cell transcriptomic analysis to study retina cell type population in treated and untreated retinas.***

Representation of data from both NHP1 and NHP2. (**A**) Integrated UMAP of the entire cell population with labelled cell types. (**B**) Major marker genes used for cell type annotation. (**C**) Proportion of cell type per AAV vector injected blebs. No major differences observed between all studied blebs. AC = amacrine cells; BPC = bipolar cells; HC = horizontal cells; IC = immune cells; MG = Müller glia; RPE = retinal pigment epithelium.

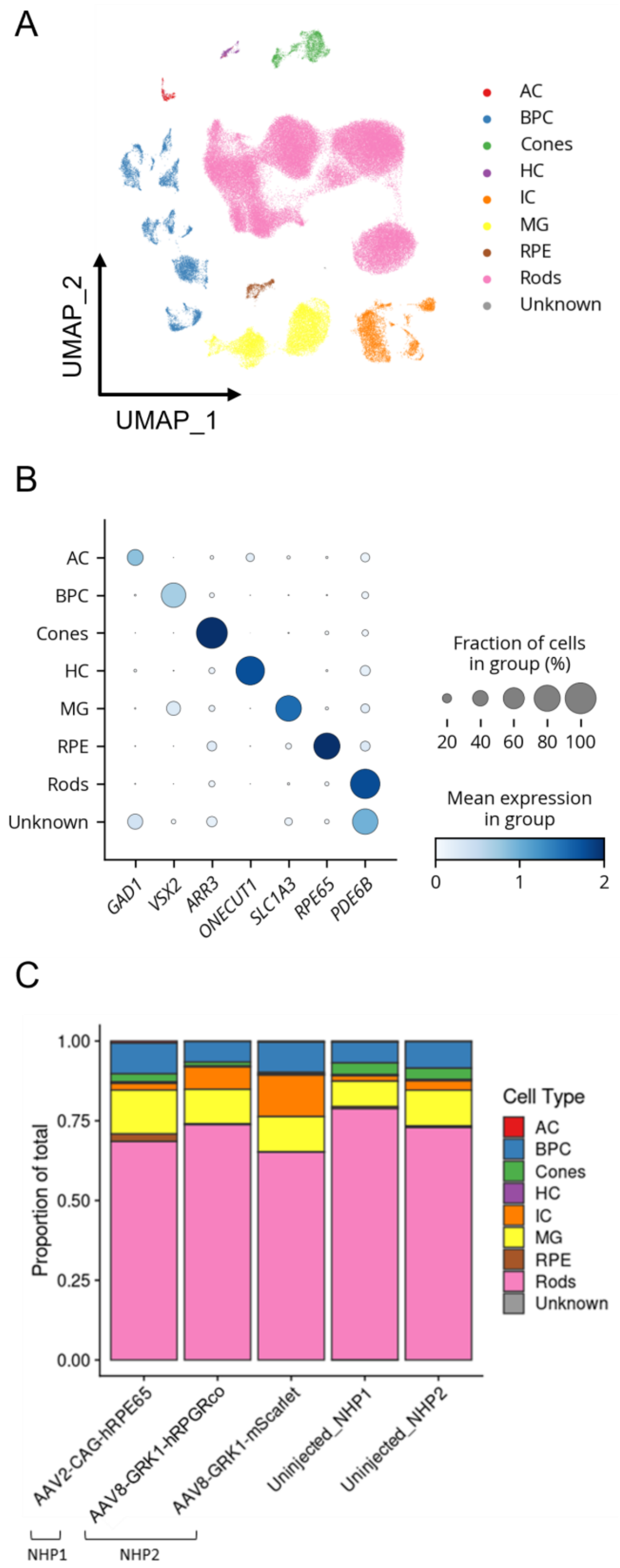

***Figure S5: No connection between apoptotic marker expression and transgene expression in the retina of AAV treated NHPs.***

Analysis of apoptotic marker expression using the Apoptosis MSigDB Hallmark gene set in rods, cones, Müller glia and RPE split by sample.

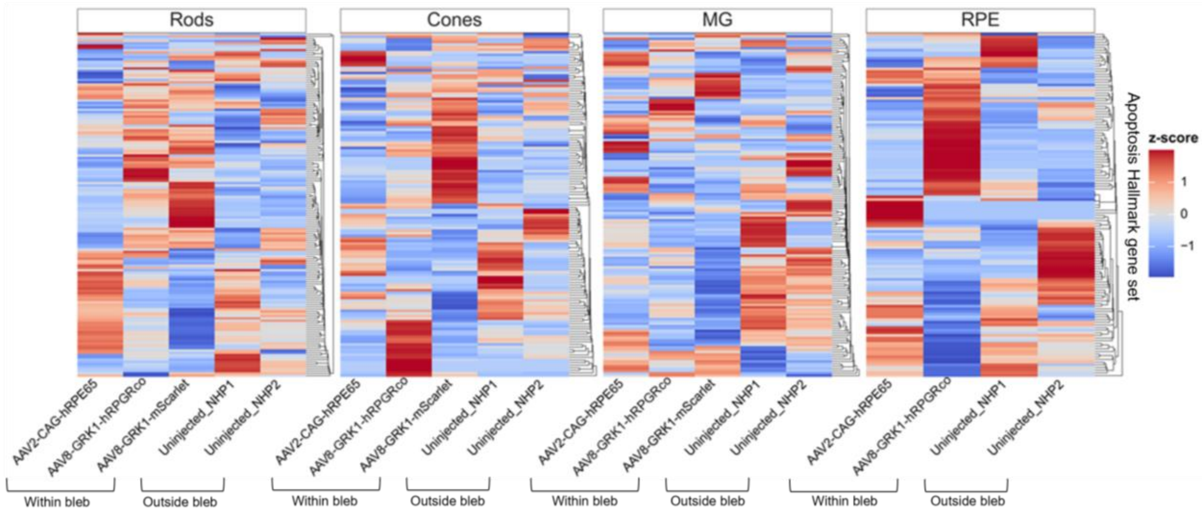

***Figure S6: Analysis of inflammation in the AAV-injected NHP retinas.***
(**A**) Immunostaining of retina sections from AAV2-CAG-*hRPE65* (NHP1) against GFAP and CD45 proteins. Control sections were taken from outside the treated blebs. Scale bars = 20µm. GCL = ganglion cell layer; INL = inner nuclear layer; OPL = outer plexiform layer; ONL = outer nuclear layer; PL = photoreceptor layer; RPE = retinal pigmented epithelium. (**B**) Spatial transcriptomic maps of B cell, T cell and natural killer (NK) cell clusters within retina sections. Coloured spots represent the locations of cells expressing the genes of interest overlayed on the H&E staining image. Scale bar = 0.5 mm.

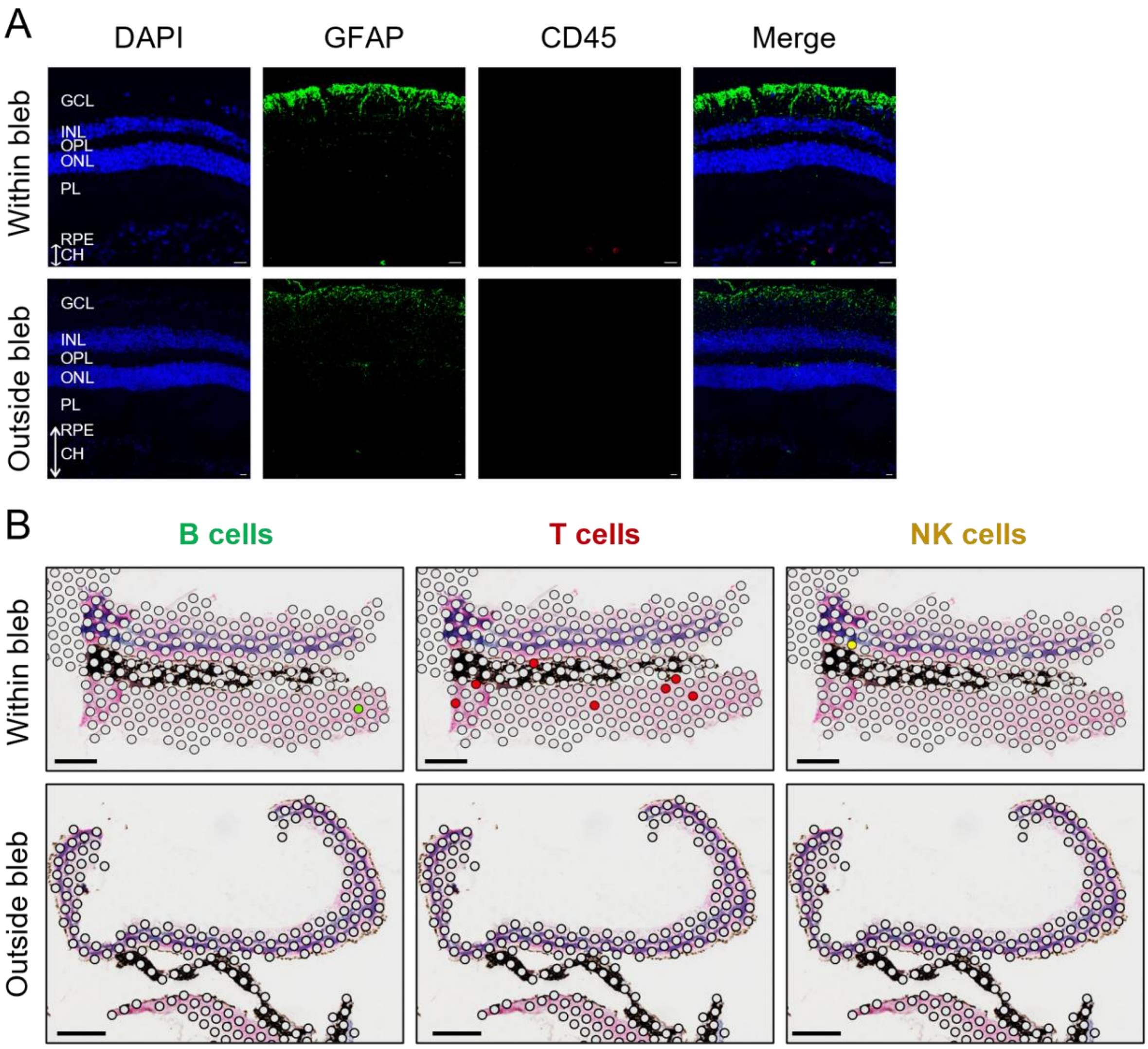

(**C**) Immunostaining of retina sections from AAV2-CAG-*hRPGRco* (NHP2) against GFAP and CD45 proteins, showing signs of inflammation. Control sections were taken from outside the treated blebs. Scale bars = 20µm. GCL = ganglion cell layer; INL = inner nuclear layer; OPL = outer plexiform layer; ONL = outer nuclear layer; PL = photoreceptor layer; RPE = retinal pigmented epithelium. (**C**) Spatial transcriptomic maps of B cell, T cell and natural killer (NK) cell clusters within retina sections. Coloured spots represent the locations of cells expressing the genes of interest overlayed on the H&E staining image. Scale bar = 0.5 mm.

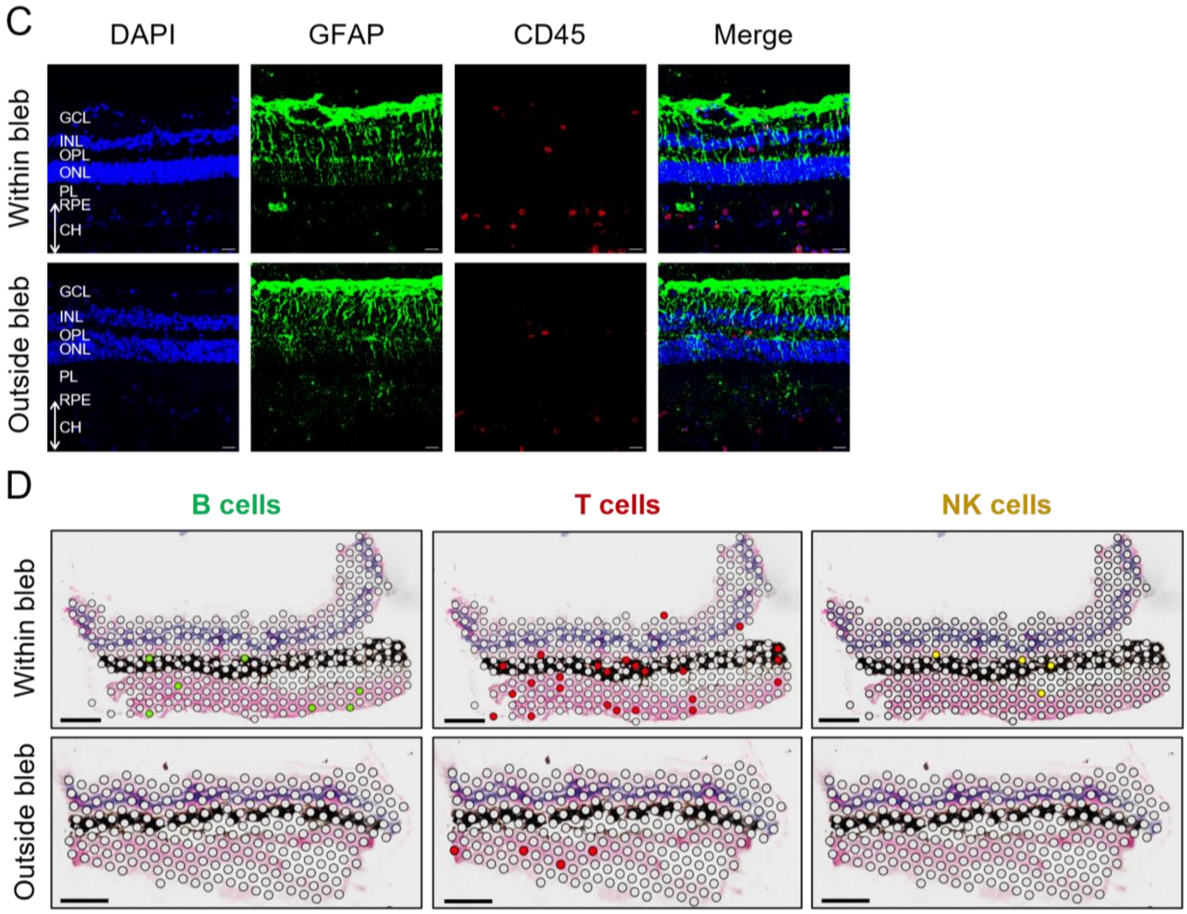

***Figure S7: Upregulated antiviral and MHC Class I genes in NHP photoreceptors after AAV gene therapy.***

Representation of data from NHP2. (**A**) DESeq2 differential expression analysis. Differentially upregulated genes include B2M (MHC I light chain), ENSMMUG00000054038 (macaque MHC I antigen), ENSMMUG00000064120 (ortholog to CD1D, MHC-like lipid antigen presenter), ENSMMUG00000050829 (MHC I pathway regulator), ENSMMUG00000052293 (ortholog to KIAA1109, MHC I complex assembly) and ENSMMUG00000058325 (MHC I antigen), indicating upregulated MHC class I antigen presentation. Upregulation of multiple immune system-associated genes in rods and cones from AAV8-GRK1-*mScarlet*-treated bleb versus retina sections outside of treated bleb. (**B**) Gene Ontology (GO) Biological Process enrichment of upregulated genes in rods. A variety of antiviral and other immune-related gene sets in AAV8-GRK1-mScarlet treated blebs can be observed when compared to retina sections outside of treated bleb. (**C**) Venn diagram comparing photoreceptors upregulated genes between AAV8-GRK1-*hRPGRco* and AAV8-GRK1-*mScarlet* treated blebs. A majority of upregulated genes in rods from AAV8-GRK1-*hRPGRco* treated blebs overlaps with those from AAV8-GRK1-*mScarlet*, suggesting a similar response. DE genes = differently expressed genes.

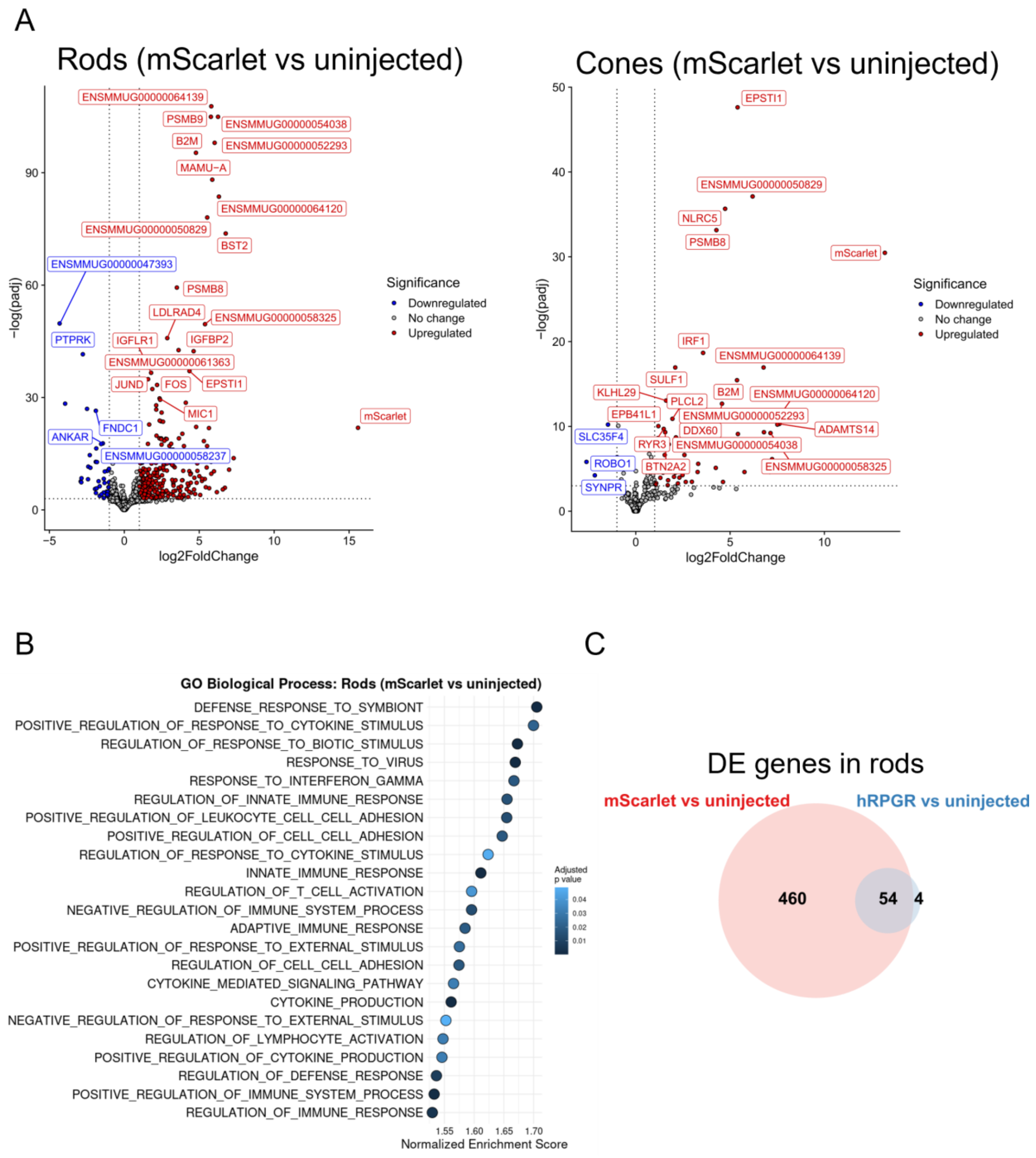

***Figure S8: The immune cell population in AAV-injected NHP retinas.***

See Figure 5 A. (**A**) Genes used to identify B cells (*MS4A1*), Perivascular cells (*RGS5*), Myeloid cells (*ITGAM*) and T cells (*CD3D*). Representation of data from both NHP1 and NHP2.

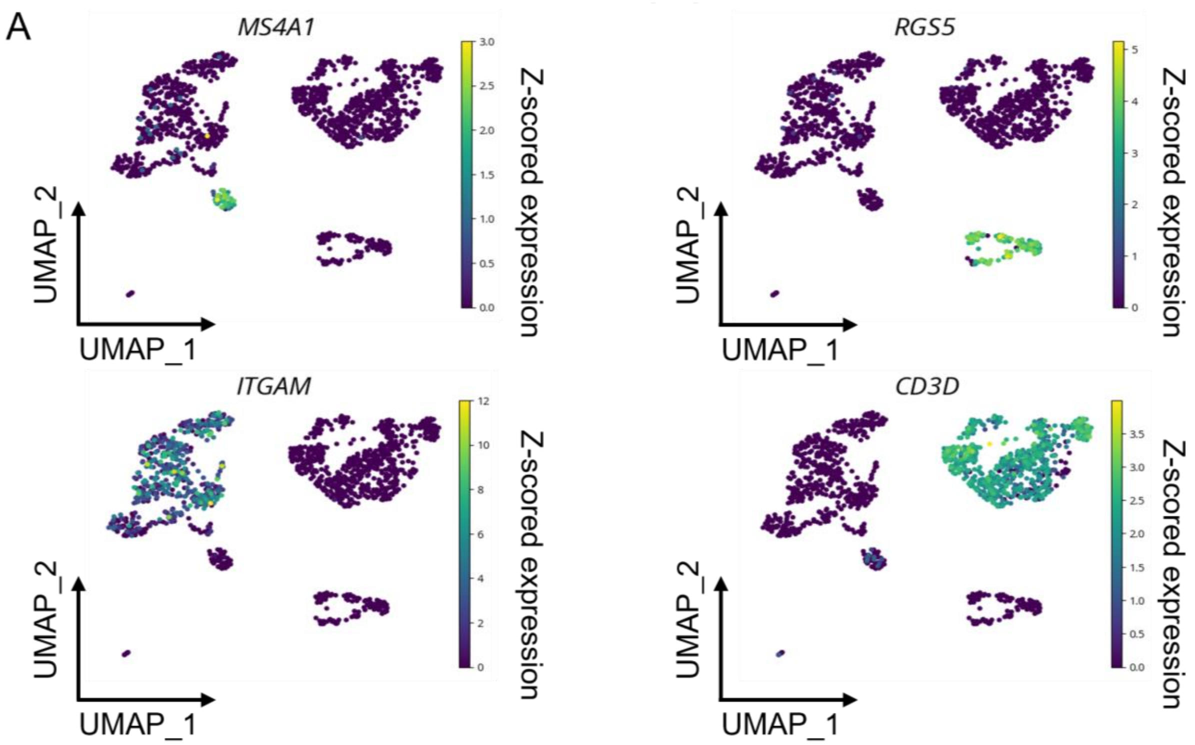

(**B**) Proportion of all immune cells separated by cell type. Predominant myeloid and T cell infiltrate. (**C**) Proportion of all immune cells per retina sections. AAV8-GRK1-*mScarlet* and AAV8-GRK1-*hRPGRco* injected blebs contributed the most to the immune cells analysed. (**D**) Proportion of each cell type per retina sections. The majority of B cells, T cells and myeloid cells came from AAV8-GRK1-*mScarlet* and AAV8-GRK1-*hRPGRco*, indicating an ongoing adaptive immune response in NHP2's retina.

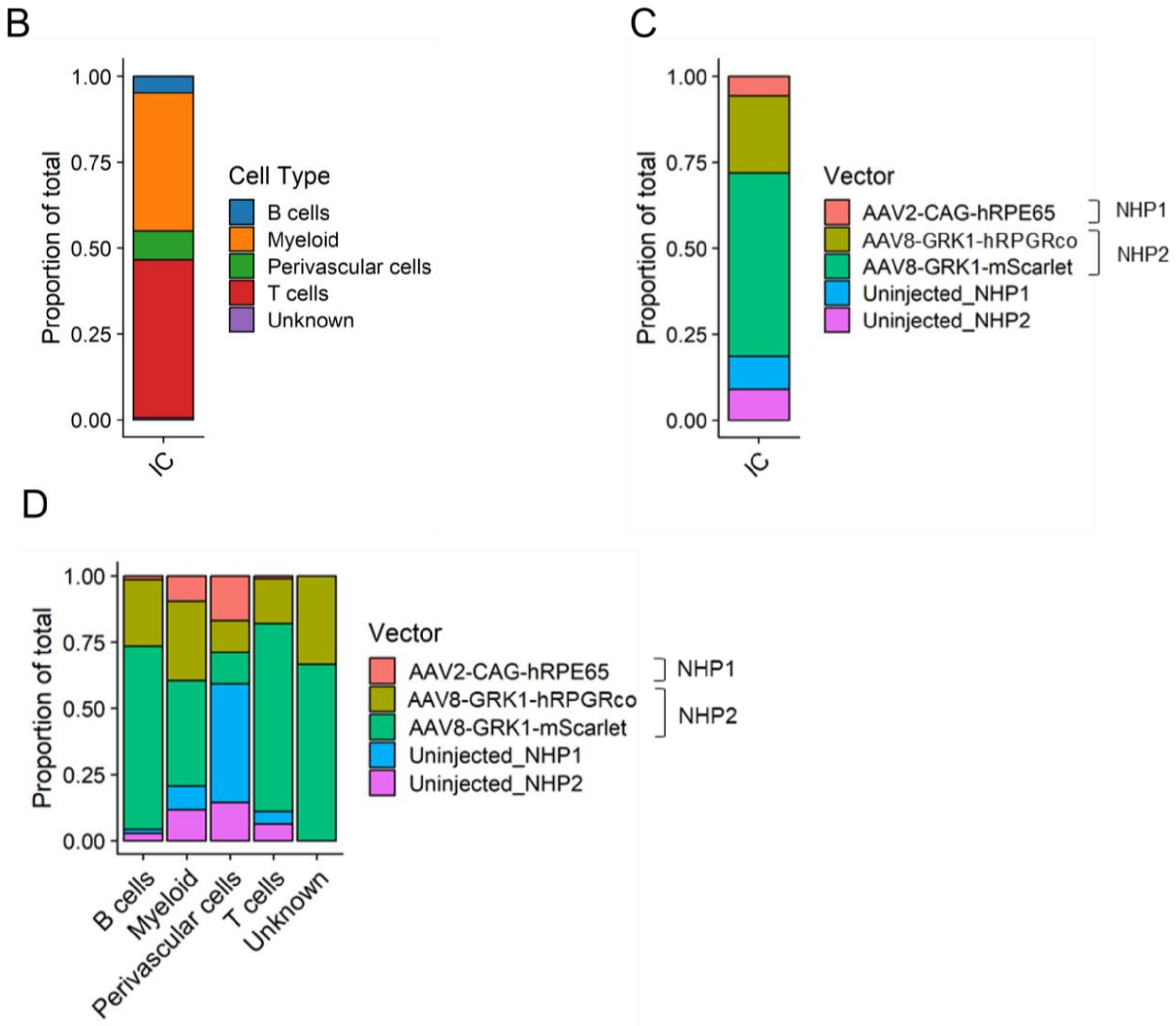

***Figure S9: Characterization of the myeloid cell population in AAV-injected NHP retinas using single-cell transcriptomic analysis.***

See Figure 5 B-C. Representation of data from both NHP1 and NHP2. (**A**) 2-dimensional FR force-directed graph of myeloid cells coloured by blebs. (**B**) Expression of microglia homeostatic markers *P2RY12* and *CX3CR1* identified the root for pseudotime analysis.

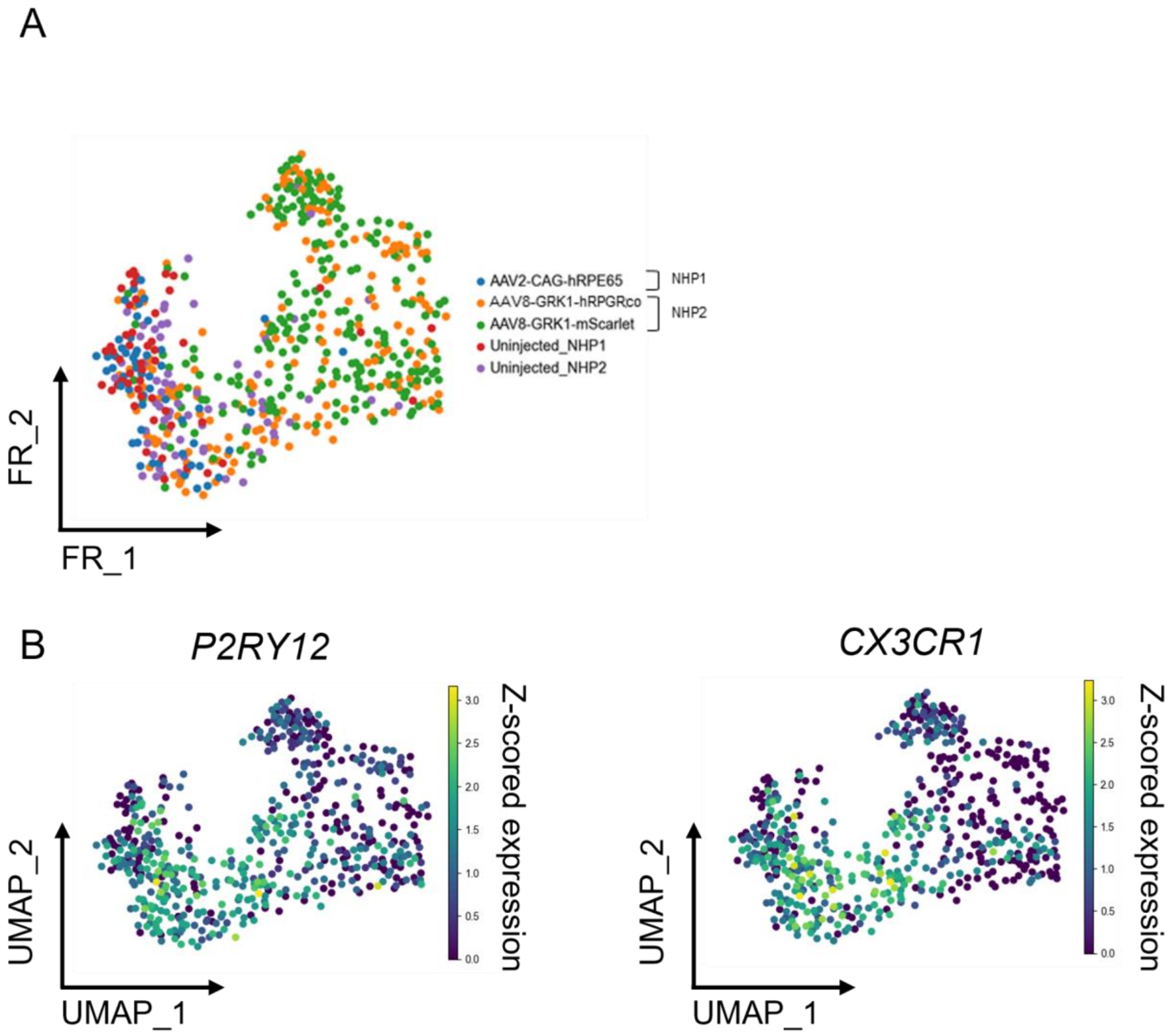

(**C**) Myeloid cells coloured by all cycle phase. No obvious proliferating cell cluster detected. (**D**) Heatmap of significant Branch 1-specific gene expressions over pseudotime. Detected MHC class I genes were previously observed in Figure S7 A. Detected MHC Class II genes included *Mamu-DRB1* (MHC II beta chain), *Mamu-DRA* (MHC class II alpha chain), *Mamu-DMB* (MHC class II peptide loader), *ENSMMUG00000056183* (ortholog to *Mamu-DQB1*, MHC II DQ beta chain) and *ENSMMUG00000019371* (ortholog to *Mamu-DQA1*, MHC II DQ alpha chain). Detected genes involved in antiviral defence included *RNASE6* (degrades viral RNA), *FGL2* (suppresses immune response, can dampen viral immune response), *BST2* (blocks virus release), *IFI27* (inhibits viral replication), *IFI6* (prevents virus-induced apoptosis), *NPC2* (regulates lipid homeostasis, affecting viral entry and replication) and *STING1* (activates interferon response). Genes commonly associated with pro-inflammatory myeloid cells such as *APOE* (inflammation marker), *APOC1* (modules immune response), *LYZ* (Bacterial defence enzyme), *FCER1G* (activates immune cells) and *C1QB* (complement system component) were also upregulated in Branch 1. (**E**) Heatmap of significant Branch 2-specific gene expressions over pseudotime. Upregulated genes were *MAP4K4* (regulates cell migration), *NR4A3* (transcription factor, migration), *MERTK* (phagocytosis and mobility), *MYO1E* (actin-based cell movement), *PTPN1* (tyrosine phosphatase, signalling), *PTPRJ* (regulates cell adhesion and mobility*)* and *CBLB* (ubiquitin ligase, immune signalling).

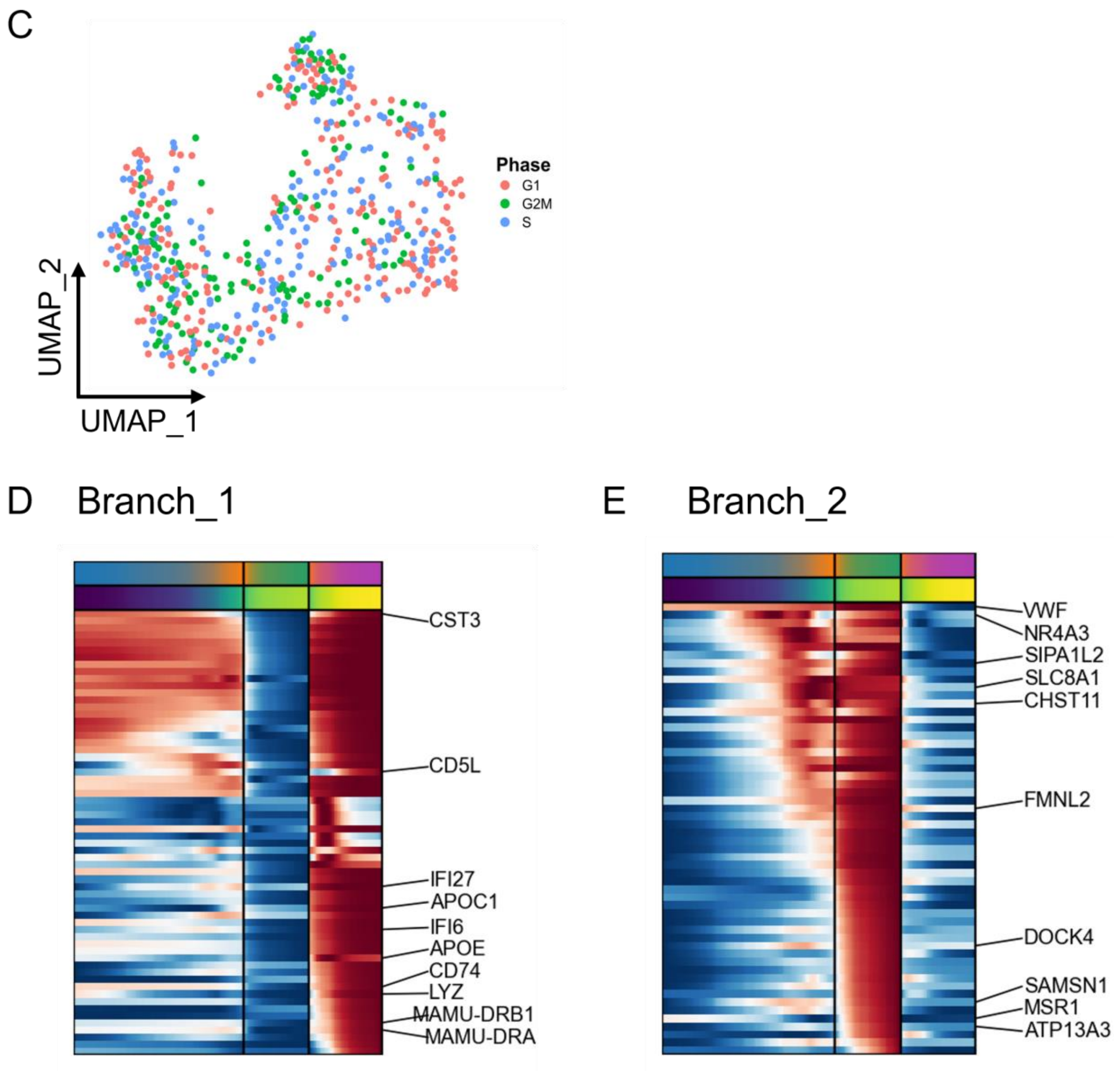

(**F**, **G**) Gene ontology (GO) enrichment analysis on Branch 1-(**F**) or Branch 2-specific genes (**G**). The majority of Branch_1 detected genes are involved in antigen presentation. Branch_2 genes are involved in cell migration and mobility, GTPase activity and protein dephosphorylation.

F Branch_1

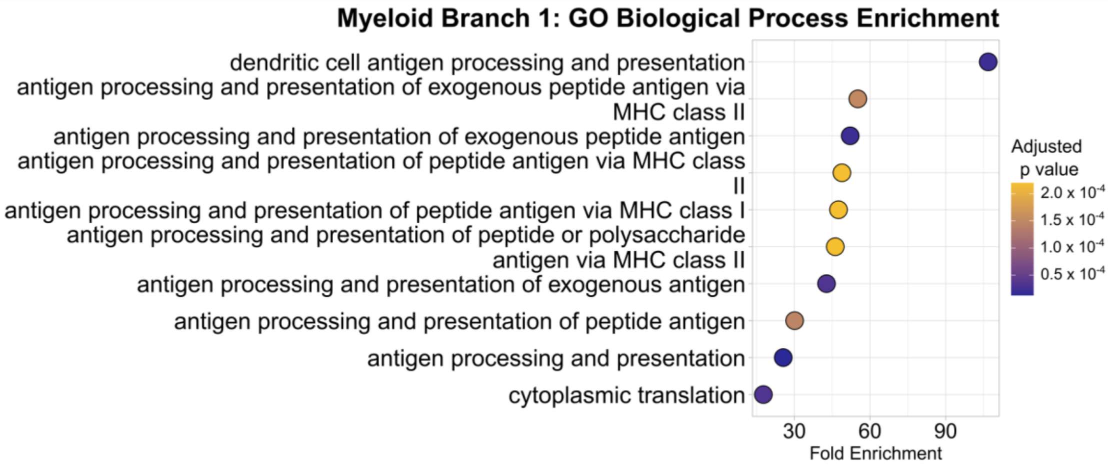

G Branch_2

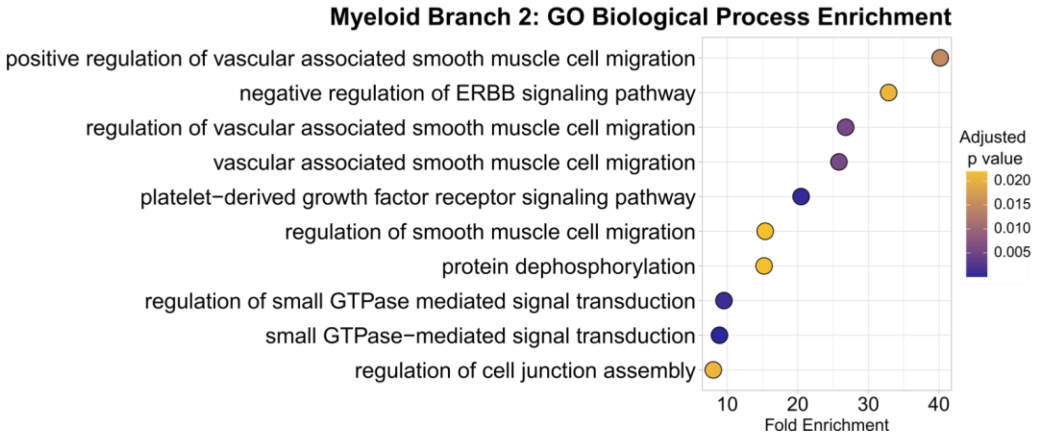

***Figure S10: Characterization of the T cell population in AAV-injected NHP retinas using single-cell transcriptomic analysis.***

See Figure 5 D-F. Representation of data from both NHP1 and NHP2. (**A**) CD4 and CD8A marker gene expressions allow differentiation between CD4+ and CD8+ T cells. (**B**) Cells coloured by cell cycle phase, highlighting a significant proportion of proliferating T cells (in S phase). (**C**) Heatmap of major T cell subsets detected based on gene expression profiles.

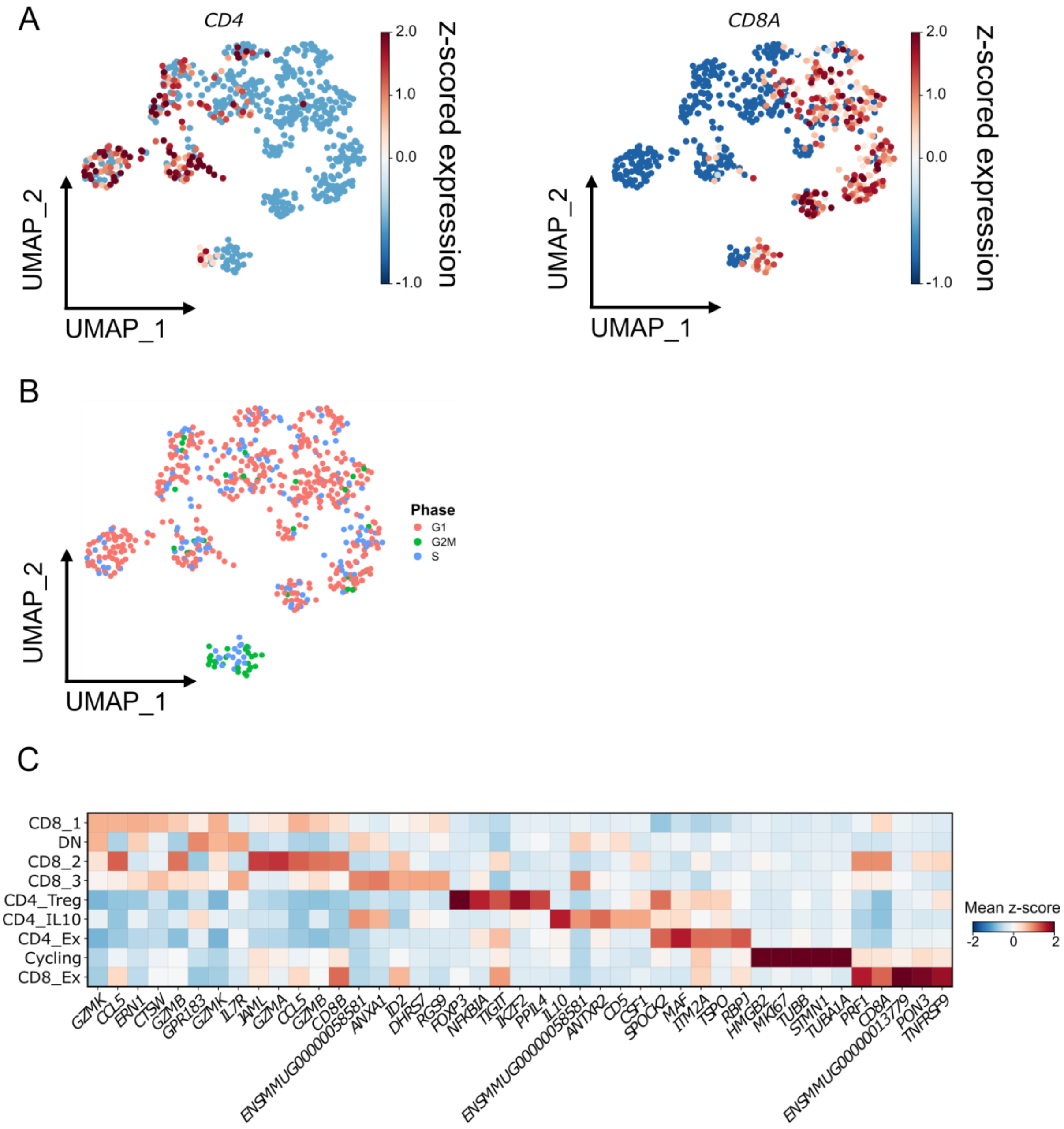

(**D**) Violin plots depicting expression of major immune checkpoint receptor genes (*PDCD1, TIGIT, LAG3 and HAVCR2*). Increased expression was seen in the 'CD8_2' T cell cluster and an exhausted CD8 T cell cluster ('CD8_Ex'). (**E**) Violin plots showing the expression of typical T follicular helper (Tfh) cell marker genes in CD4 clusters. (**F**) Scatter plots showing expression of Tfh marker genes against *CXCL13* in *CXCL13*$^{+}$ *CD4*$^{+}$ cells. Co-expression of *PDCD1*, *ICOS* and *BCL6*, but low expression of *CXCR5* might suggest a composition of T peripheral helper cells.

D

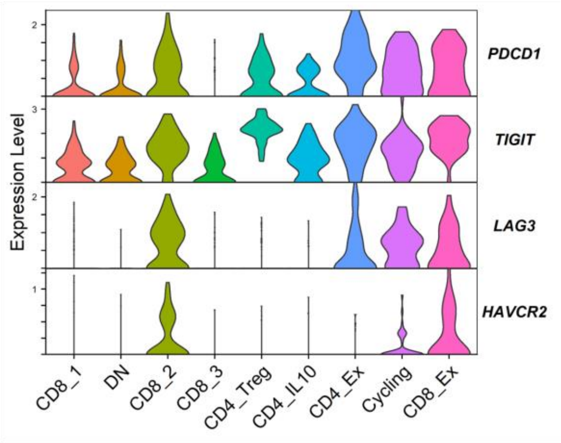

E

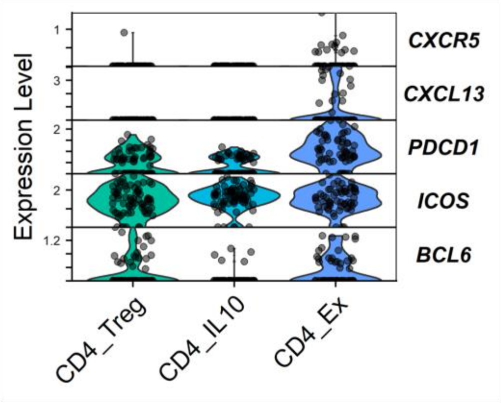

F

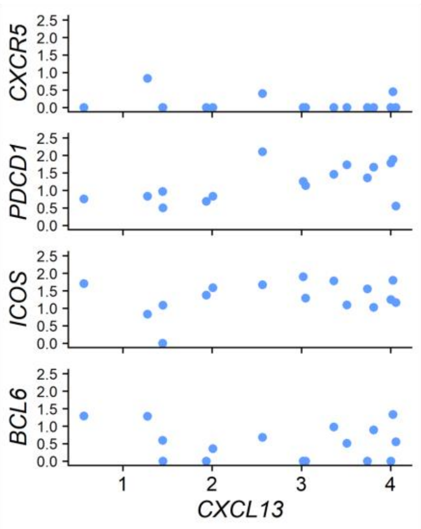

***Figure S11: Additional cytokine panels for NHP vitreous samples.***
See Figure 6. Additional cytokines assayed as part of the LegendPlex NHP Inflammation Panel in the vitreous samples from NHP1 (**A**) and NHP2 (**B**). Ratio scale is in log 2 compared to baseline.

A **NHP 1**

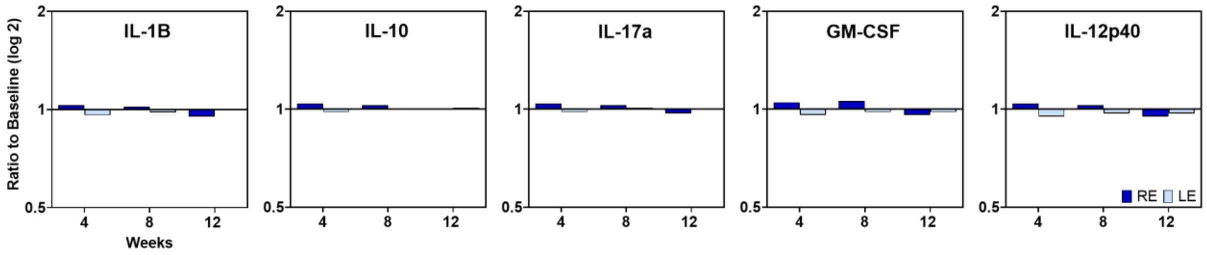

B **NHP 2**

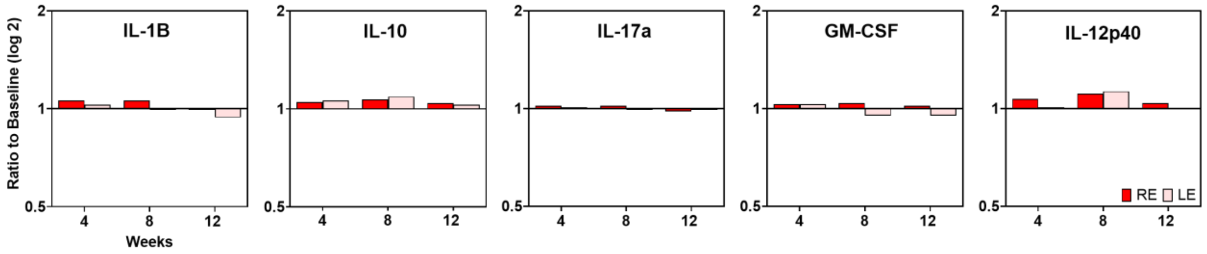

***Figure S12: Cytokine profiling of NHP peripheral blood mononuclear cells (PBMCs) revealed no major changes.***

Analysis of cytokine expression in the blood samples from NHP1 (**A**) and NHP2 (**B**) following subretinal AAV gene therapy. Ratio scale is in log 2 compared to baseline.

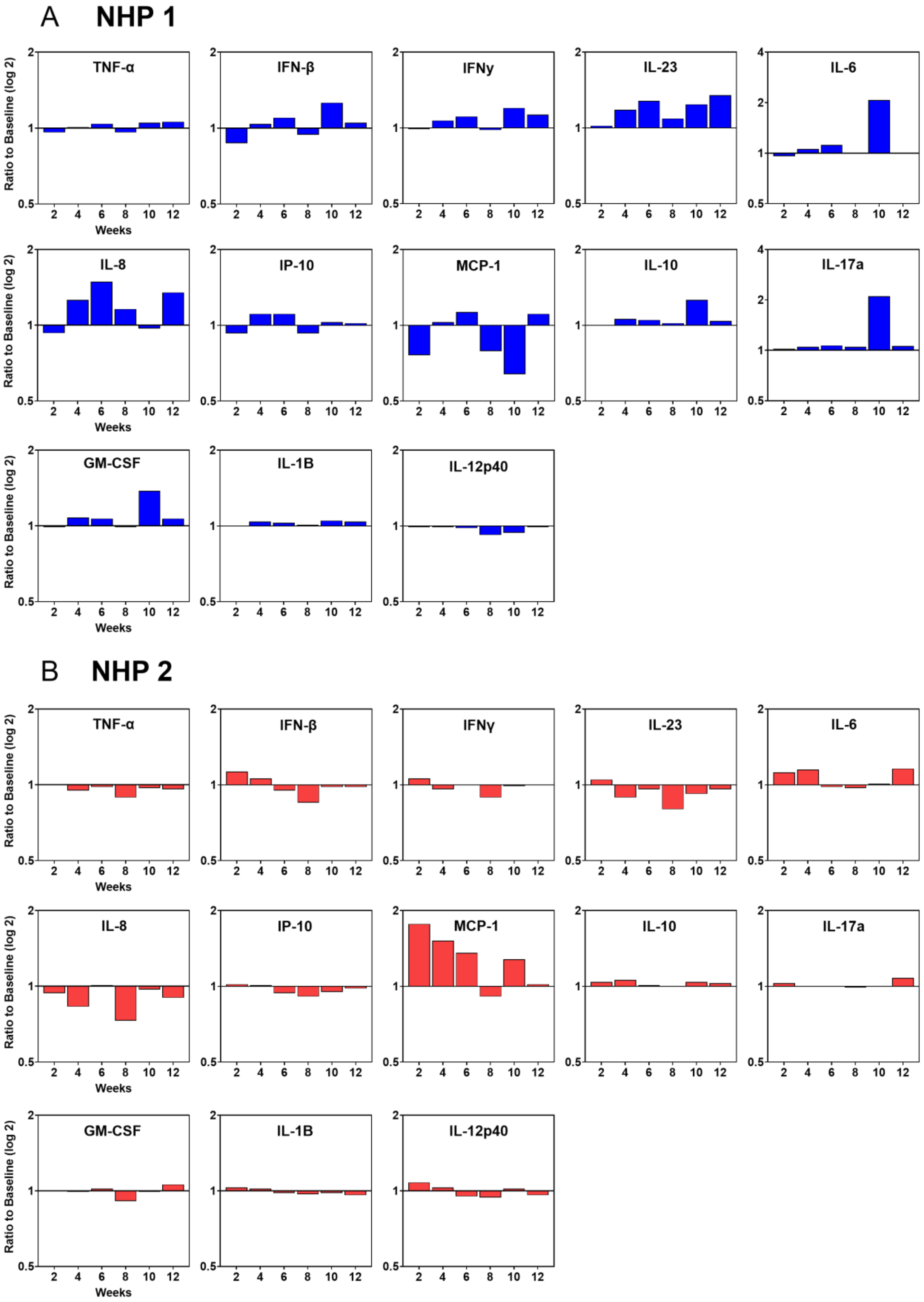

***Figure S13: Additional cytokine panel assay results from human iPSC-derived microglia.***
See Figure 7. Rest of the cytokine panel expression from human iPSC-derived microglia after AAV stimulation. Data are represented as mean ± SD. n=4. A two-way ANOVA statistical test was performed. * = vs AAV Only. * = p>0.01. ** = 0.001<p<0.01. *** = 0.0001<p<0.001.

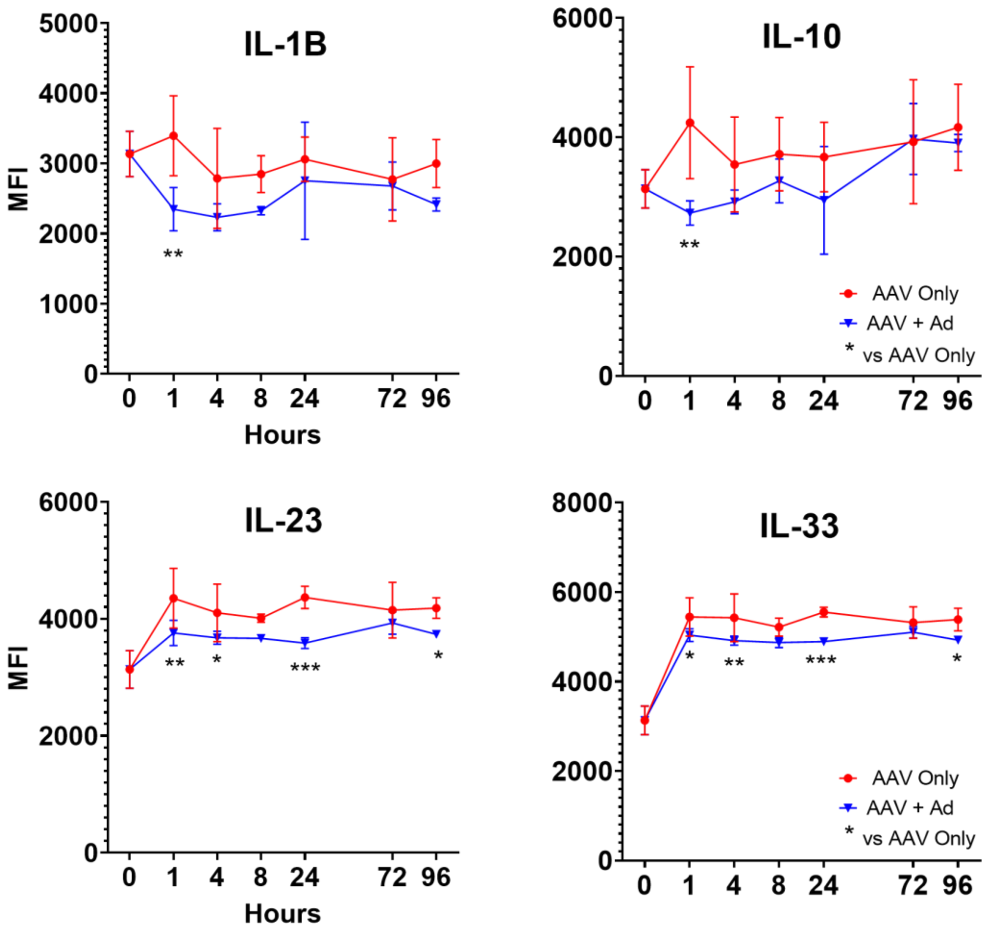

*Table S1: List of PCR primers.*

| Name | Sequence | Purpose |
|---|---|---|
| **mScarlet_F** | GCGTGATGAACTTCGAGGAC | qPCR Titration of Vector |
| **mScarlet_R** | CTTGTAGATCAGGGTGCCGT | qPCR Titration of Vector |

*Table S2: List of antibodies.*

| Target | Host | Clonality | Reference | Supplier | Working dilution |
|---|---|---|---|---|---|
| **RPGR** | Rabbit | Polyclonal | HPA001593 | Sigma-Aldrich | 1:200 (IHC) |
| **RPE65** | Mouse | Monoclonal | 401.8B11.3D9 | Novus Biologicals | 1:250 (IHC) |
| **IBA1** | Rabbit | Monoclonal | 019-19741 | Wako | 1:500 (IHC) |
| **GFAP** | Chicken | Polyclonal | Ab4674 | Abcam | 1:200 (IHC) |
| **IBA1** | Rabbit | Monoclonal | 019-19741 | Wako | 1:500 (IHC) |
| **CD45** | Rabbit | Monoclonal | Ab281586 | Abcam | 1:250 (IHC) |
